# A radial glial regulatory program bridges progenitor maintenance and neuronal identity

**DOI:** 10.64898/2026.11.16.767040

**Authors:** Jaime Fabra-Beser, Jordi Planells, Luuli Tran, Dmytro Tsurkan, Alba Marín-Garnes, Laura Veintimilla-Escot, Carmen Mateos-Martínez, Tyler Gibson, Santos J. Franco, Cristina Gil-Sanz

## Abstract

The cerebral cortex acquires its neuronal diversity through successive waves of neurogenesis. How molecular differences among early progenitors relate to their neuronal output remains unclear. Here, we identify a molecular program in early SOX9-high radial glia (S9RGs) that links progenitor maintenance with the regulation of genes associated with neuronal identity. Combining transcriptional and chromatin profiling with in vivo perturbations, we identify a program involving extracellular signaling, cell-cycle regulation, progenitor maintenance and neurogenesis. We show that SOX9 regulates part of this program, including receptors that contribute to progenitor maintenance, while Notch acts upstream of Sox9 and reproduces several transcriptional and chromatin features of S9RGs. S9RGs express selected later-stage radial glial features and show distinct chromatin accessibility patterns at genes involved in progenitor maintenance, neurogenesis and neuronal identity. Notch activation reproduces part of these changes, including differences at genes associated with lower- and upper-layer neuronal specification. Interestingly, S9RGs, which preferentially generate upper-layer neurons, share molecular features with the highly proliferative RGα population in ferrets and with human outer RGs (oRGs) from cortical organoids, despite their different positions within the developing cortex. Together, our findings suggest that molecular differences among early radial glia could influence both how long they remain as progenitors and the neuronal identities they subsequently generate.

## Introduction

The different neuronal subtypes of the cerebral cortex are generated from radial glial cells (RGs), either directly or through intermediate progenitors. These neurons are produced in a temporal sequence, with early-born neurons mainly occupying the deep layers and later-born neurons populating the upper layers. Changes in progenitor gene expression contribute to this progression and are transmitted to their neuronal descendants (1, 2). However, RGs present at the same embryonic age can differ in their neurogenic behavior and in the neurons they generate. Recent clonal analyses also suggest that differences in neuronal projection identity can arise early within RG lineages (3). How progenitors are maintained for later neurogenesis, and how this relates to the types of neurons they generate, is still poorly understood.

The contribution of developmental time and progenitor heterogeneity to neuronal specification has remained controversial. Heterochronic transplantation experiments showed that progenitor competence changes with age (4, 5). Genetic fate mapping subsequently identified early progenitor populations with different contributions to neuronal subtypes, although other lineage studies found broader laminar contributions and considerable variation among individual clones (6–10). Moreover, temporal identity can progress independently of cell division, and apical progenitors can still respond to a younger developmental environment (11, 12). These findings suggest that neuronal production depends both on changes over time and on molecular differences among progenitors. However, we still do not know whether progenitors maintained for later neurogenesis simply delay their differentiation or whether their molecular properties also influence how they generate particular neuronal identities.

SOX9 regulates neural stem cell specification and maintenance (13). Moreover, it can promote extracellular matrix production and basal progenitor proliferation during neocortical development (14). In our previous work, we identified a population of early Hes5-reporter-positive RGs with high Sox9 expression that maintain their progenitor state for longer, delay differentiation and preferentially generate upper-layer neurons. We also showed that modifying Sox9 levels changes progenitor maintenance, cell-cycle progression and the neurons subsequently generated (15). However, the molecular program associated with these differences in endogenous SOX9 expression, its regulation and its potential contribution to neuronal specification remained unknown.

Notch signaling could participate in the regulation of these processes. It maintains RGs and regulates their differentiation into intermediate progenitors and neurons through changes in the activity of its downstream effectors (16–19). In addition, Notch has been implicated in neuronal specification and, at later stages, in chromatin changes that allow progenitors to generate glial cells (20, 21). Interestingly, genes associated with neuronal identity are already expressed in cortical progenitors before differentiation. Moreover, altering chromatin regulators in progenitors affects the neurons they later generate (22–24). These observations raise the question of whether signals that maintain early RGs also contribute to the regulation of neuronal programs before differentiation.

Here, we use a Sox9 promoter reporter to isolate early SOX9-high RGs and study their gene expression and chromatin organization. We show that these cells express several receptors and share molecular features with later progenitors. We demonstrate that Sox9 regulates part of this receptor program, that selected receptors contribute to progenitor maintenance and that Notch regulates Sox9 expression. We also identify differences in chromatin accessibility at genes associated with neuronal identity and find components of the S9RG molecular program in progenitors from other mammalian species. Our results suggest that, in addition to delaying neurogenesis, these cells have early molecular differences that could contribute to the neurons they later generate.

## Results

### Sox9 promoter activity identifies early radial glial cells with delayed neurogenesis

We previously showed that early progenitors with higher SOX9 expression delay neurogenesis (15). To identify these progenitors and study their molecular properties and behavior, we generated a construct expressing nuclear GFP under the control of a proximal fragment of the mouse Sox9 promoter (Sox9-nGFP). We electroporated this construct together with CAG-mCherry into the mouse cortex at E12.5 and analyzed the labeled cells at E13.5 (Fig. 1A–C). Most cells expressing Sox9-nGFP were located in the ventricular zone (VZ). In contrast, cells labeled with the constitutive reporter were distributed across the cortical wall, including regions containing differentiating cells (Fig. 1B–C). Moreover, we confirmed by immunostaining that Sox9-nGFP-positive cells in the VZ expressed higher levels of endogenous SOX9 than reporter-negative RGs (Fig. 1D).

**Figure 1.**
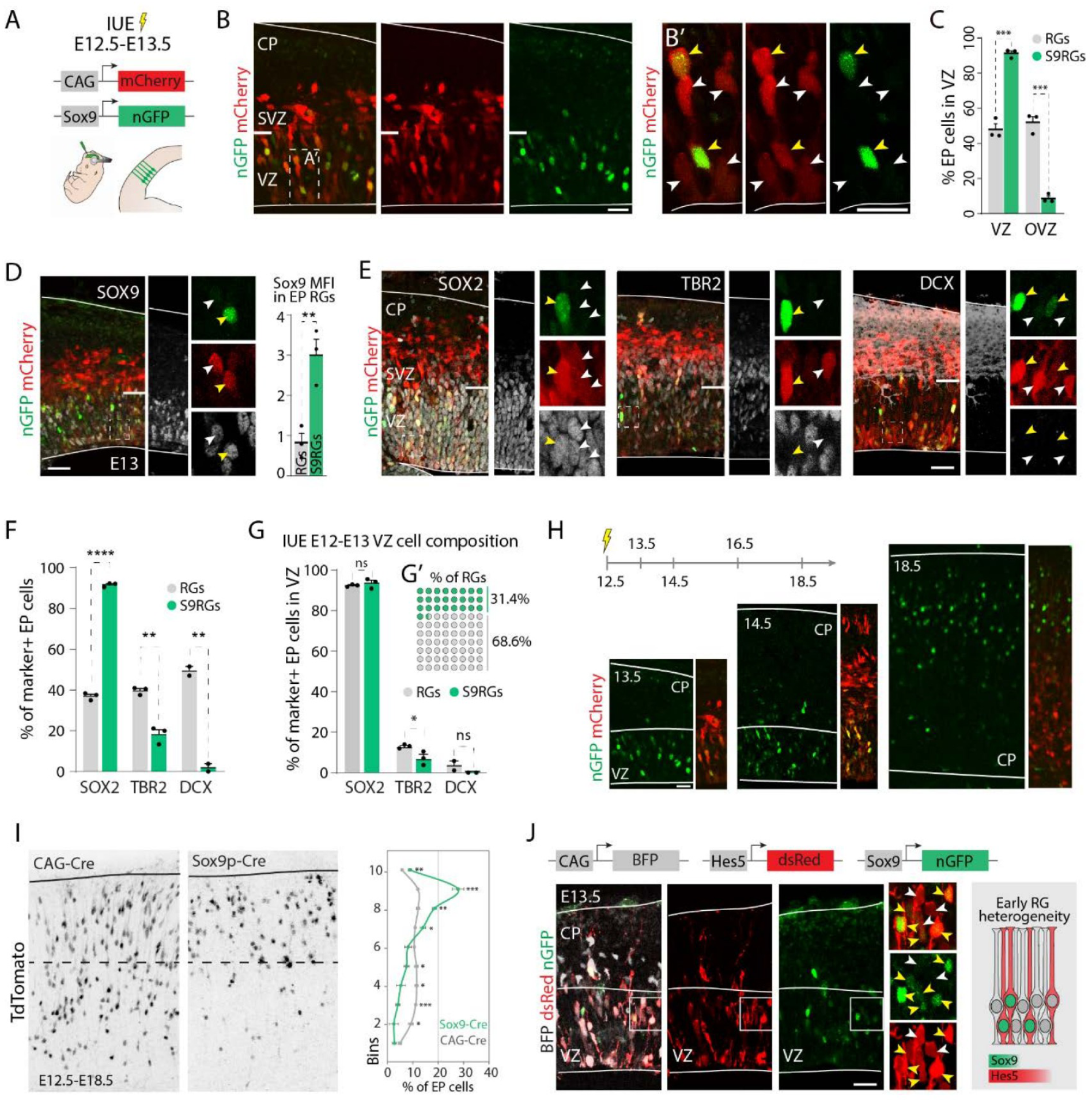
Sox9 promoter activity enables prospective targeting of early radial glia with delayed neurogenesis. A, Co-electroporation of CAG-mCherry and *Sox9* promoter-nuclear GFP at E12.5, with collection at E13.5. B, Reporter distributions and ventricular-zone magnification. C, Distribution of labeled cells inside and outside the ventricular zone. D, Endogenous SOX9 immunostaining and mean fluorescence intensity in reporter-positive and reporter-negative ventricular-zone cells. E-G, SOX2, TBR2 and DCX marker analysis, including the composition of the ventricular-zone population and the reporter-positive fraction. H, Distribution of labeled cells at E13.5, E14.5 and E18.5. I, Ai9 labeling after CAG-Cre or *Sox9*-Cre electroporation at E12.5, analyzed at E18.5; the cortical plate was divided into ten bins. J, Co-labeling with CAG-BFP, *Hes5*-dsRed and *Sox9*-nGFP. Graphs show mean and SEM where indicated. VZ, ventricular zone; CP, cortical plate; EP, electroporated. Related supplementary figure: S1.

We examined whether the reporter distinguished RGs or simply labeled different proportions of progenitors and differentiating cells. Across the cortical wall, a higher proportion of reporter-positive cells expressed SOX2, while fewer expressed the intermediate progenitor marker TBR2 or the neuronal marker DCX (Fig. 1E–F). However, within the VZ, more than 90% of both reporter-positive and reporter-negative electroporated cells expressed SOX2. Cells expressing the Sox9 reporter represented approximately 30% of the electroporated SOX2-positive RGs (Fig. 1G). These results show that the reporter distinguishes two populations of RGs with different levels of SOX9. We refer to reporter-positive progenitors as S9RGs and reporter-negative progenitors as comparison RGs.

We followed the labeled cells at later ages after electroporation. Cells labeled with the Sox9 reporter remained in the VZ for longer, while more cells labeled with the constitutive reporter were already located in differentiating regions (Fig. 1H). Moreover, using Sox9-promoter-driven Cre in Ai9 embryos, we found that their descendants were preferentially located in the upper cortical layers at E18.5 (Fig. 1I). These results indicate that cells with early Sox9 promoter activity delay neurogenesis and preferentially contribute to later-born neurons. We also examined the relationship between S9RGs and the Hes5-positive progenitors described in our previous work (15). Both reporters showed considerable overlap, although some Hes5-positive cells did not show detectable Sox9 reporter activity (Fig. 1J).

Having established that the Sox9 reporter identifies a population of RGs with delayed neurogenesis, we sought to characterize their molecular properties. However, the labeled populations contained different proportions of differentiated cells, so we first isolated RGs to avoid differences in cell composition. For this, we used CD133 and CD24 to select viable CD133-positive cells with low or no CD24 expression, and then separated them according to Sox9 reporter expression. Intracellular staining for SOX9, TBR2 and DCX allowed us to distinguish progenitors from differentiating cells. Moreover, qPCR confirmed that both sorted populations were enriched in RG markers compared with neurons (fig. S1). This way, we could compare RGs with different Sox9 reporter activity while reducing differences due to the presence of differentiating cells.

### Early SOX9-high progenitors share molecular features with later RGs

To identify molecular differences associated with delayed neurogenesis in S9RGs, we performed RNA sequencing of four independently sorted samples per population, obtained from three litters. Principal component analysis separated S9RGs from comparison RGs (Fig. 2A–B). We identified 310 differentially expressed genes, of which 103 were increased and 207 were reduced in S9RGs (adjusted P ≤ 0.05; Fig. 2C–D). Moreover, markers of RGs, intermediate progenitors and neurons did not show consistent differences suggesting that one population contained more differentiated cells, in agreement with our sorting validations.

**Figure 2.**
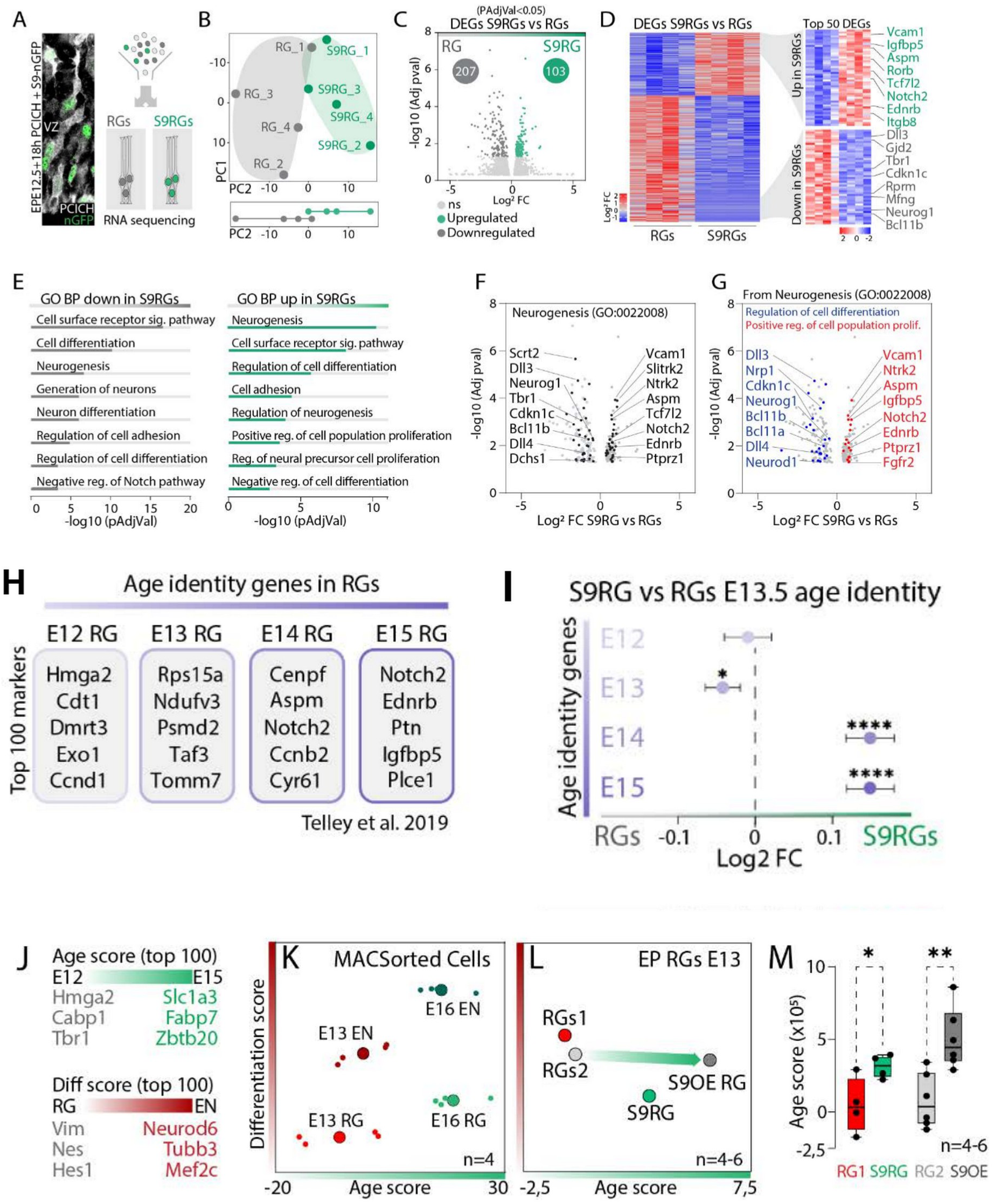
Early SOX9-high progenitors engage a later radial glial program. A, Isolation of reporter-defined RGs after E12.5 electroporation. B, Principal component analysis of four independently sorted biological replicates per condition. C-D, Differential expression and heatmaps, with 103 enriched and 207 reduced genes in S9RGs at adjusted P ≤ 0.05. E-G, Functional enrichment and neurogenesis-associated differential genes. H-I, Reference temporal RG gene sets and their expression in the age-matched S9RG comparison, using Telley et al. (1). J, Reference age and differentiation scores from the same study. K, Score validation in independently isolated E13 and E16 CD133-enriched RG and neuron-enriched fractions, n = 4. L-M, Scores in endogenous S9RGs and *Sox9*-overexpressing RGs with their respective batch-matched controls, n = 4–6. Age scores indicate temporal gene-set enrichment rather than directly measured developmental age. Related supplementary figures: S1, S2.

Gene Ontology analysis associated these genes with neurogenesis, receptor signaling, adhesion and the regulation of proliferation (Fig. 2E). Genes related to neurogenesis were found among both upregulated and downregulated genes, although they included different components of this process. For example, S9RGs expressed higher levels of Notch2, Ednrb, Ptprz1, Vcam1 and Aspm, while Neurog1, Dll3 and other genes associated with neurogenic progression were reduced (Fig. 2F–G). We validated these expression differences by qPCR, confirming increased expression of selected signaling genes and reduced expression of genes associated with differentiation. Moreover, RNAscope showed higher expression of Vcam1, Notch2, Ednrb and Ptprz1 in reporter-positive cells in the VZ compared with neighboring electroporated reporter-negative cells (fig. S2). These observations confirmed the expression differences within progenitors in the tissue, rather than differences in the proportions of cell types in the sorted samples.

Interestingly, several receptors and adhesion-related genes increased in S9RGs are normally expressed at higher levels in later RGs. We compared these changes with the temporal RG gene sets described by Telley et al. (1). Although both populations were isolated at the same embryonic age, S9RGs showed lower expression of selected early RG genes and higher expression of genes associated with E14–E15 RGs (Fig. 2H–I). To determine whether these differences reflected developmental age rather than differentiation, we used the gene scores defined for each process in the same study. We first tested them in independently isolated RG- and neuron-enriched populations from E13 and E16 cortices. The age score distinguished the two developmental stages, while the differentiation score separated RGs from neurons (Fig. 2J–K). Applying these scores to S9RGs and comparison RGs isolated at the same early age revealed that S9RGs had a higher age score, without a significant increase in the differentiation score (Fig. 2L–M). Thus, the increased expression of receptors and other signaling-related genes in S9RGs is associated with later RG features without evidence of increased differentiation.

### SOX9 promotes a receptor program that restrains neuronal differentiation

To determine whether Sox9 regulates genes enriched in S9RGs, we electroporated a Sox9 expression construct or the control vector at E12.5 and isolated the labeled RGs 16–18 h later. We chose this early time point to study changes in gene expression before the differences in progenitor and neuronal distribution described in our previous work (15). RNA sequencing of six biological replicates per condition revealed 768 differentially expressed genes (Fig. 3A–B). Among them, genes involved in neurogenesis, including Neurog1 and Neurod1, were reduced, while progenitor-associated genes were increased. We confirmed selected changes by qPCR, including the increased expression of Col2a1, a gene known to respond to Sox9 (Fig. 3C–E). Comparison with the S9RG expression profile showed that Sox9 overexpression increased the expression of Notch2, Ednrb and Ptprz1 and reduced genes associated with differentiation, as we had observed in S9RGs (Fig. 3F–H). Some of these receptors reached levels similar to those found in S9RGs, indicating that Sox9 can promote the expression of part of this molecular program. Moreover, Sox9 overexpression increased the age score without a significant increase in the differentiation score (Fig. 2L–M). Together, these changes suggest that Sox9 promotes part of the later RG-associated program without increasing differentiation.

**Figure 3.**
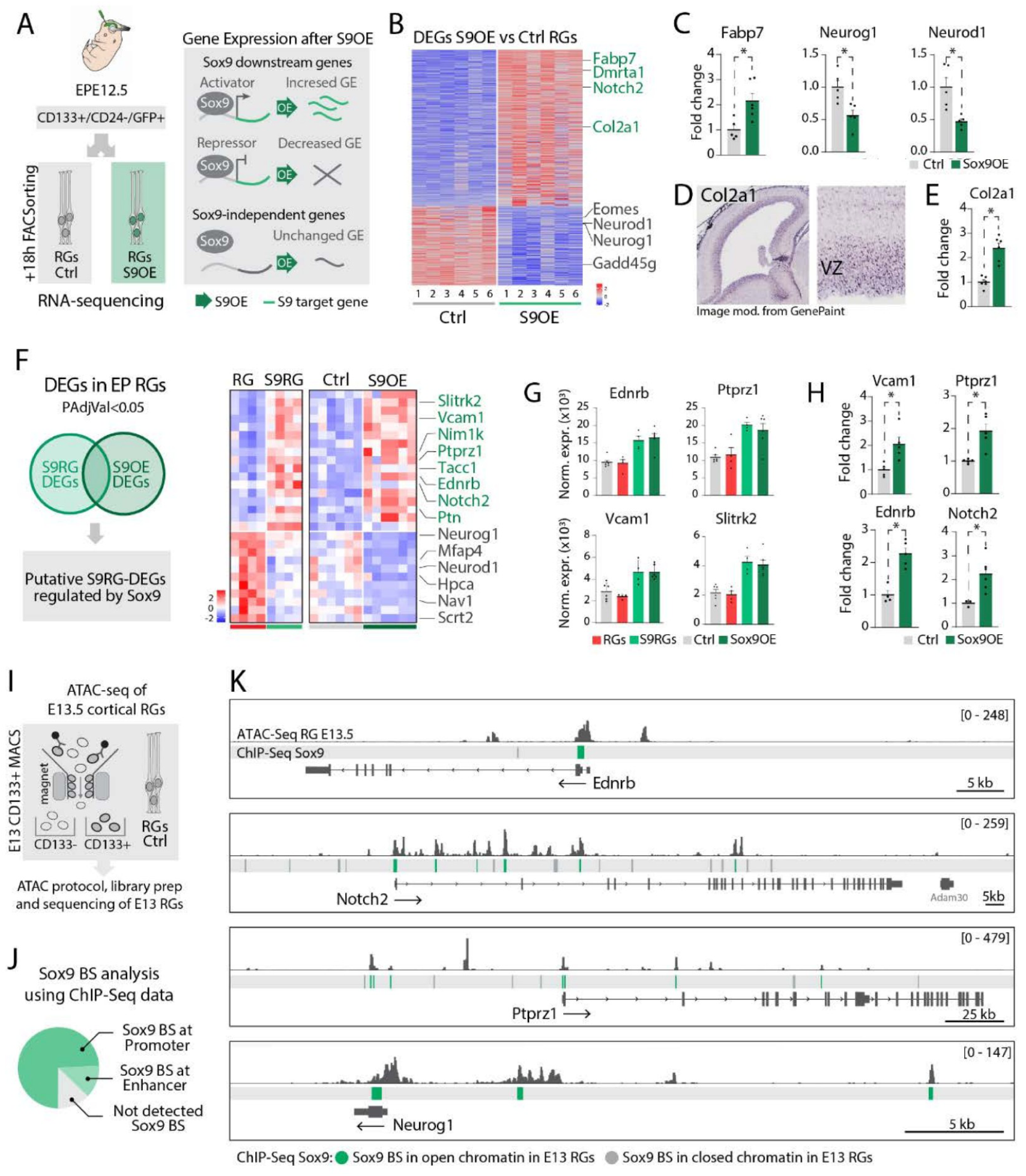
SOX9 promotes the maintenance-associated receptor program. A, Electroporation of control PCIG or PCIG-*Sox9* at E12.5 and sorting of RGs approximately 16-18 h later. B, Differential expression after *Sox9* overexpression, six biological replicates per condition. C, qPCR validation of Fabp7, *Neurog1* and *Neurod1*, five to six biological samples. D-E, *Col2a1* reference expression and qPCR response. F, Overlap of *Sox9*-responsive and endogenous S9RG differential genes. G-H, Selected normalized expression values and qPCR validations. I, CD133-based enrichment of early RGs for accessibility profiling. J-K, Overlap with public SOX9 ChIP-seq evidence and accessible regions at selected loci. Public SOX9 binding annotations identify candidate regulatory elements within accessible cortical RG chromatin. Related supplementary figure: S3.

To examine potential regulatory regions associated with these genes, we compared publicly available SOX9 ChIP-seq data with regions of accessible chromatin in early cortical RGs. We found that some of the reported SOX9 binding sites were located in accessible regions near genes whose expression changed after Sox9 overexpression, including receptors and genes involved in neurogenesis (Fig. 3I–K). These regions could therefore participate in the regulation of these genes by SOX9. To further examine the regulation of these receptors in cortical progenitors, we analyzed Notch2 and Ednrb expression by RNAscope after Sox9 overexpression or knockdown. Sox9 overexpression increased the proportion of electroporated VZ cells with high transcript levels, while Sox9 knockdown reduced this proportion (fig. S3). These results indicate that the expression of both receptors changes with Sox9 levels within cortical progenitors.

To test whether these receptors contribute to S9RG maintenance, we co-electroporated the Sox9 reporter with shRNAs against Notch2 or Ednrb at E12.5 and analyzed the cortex at E14.5 (Fig. 4A). Reducing the expression of either receptor decreased the proportion of reporter-positive cells remaining in the VZ (Fig. 4B–C). We confirmed by RNAscope that the corresponding transcript was reduced in electroporated cells (Fig. 4D–E). We also examined the identity of reporter-positive cells outside the VZ after receptor knockdown. These cells expressed NEUROD2 following either Notch2 or Ednrb knockdown (Fig. 4F–H). Together, these findings indicate that reducing Notch2 or Ednrb promotes neuronal differentiation and support a role for these Sox9-regulated receptors in maintaining S9RGs.

**Figure 4.**
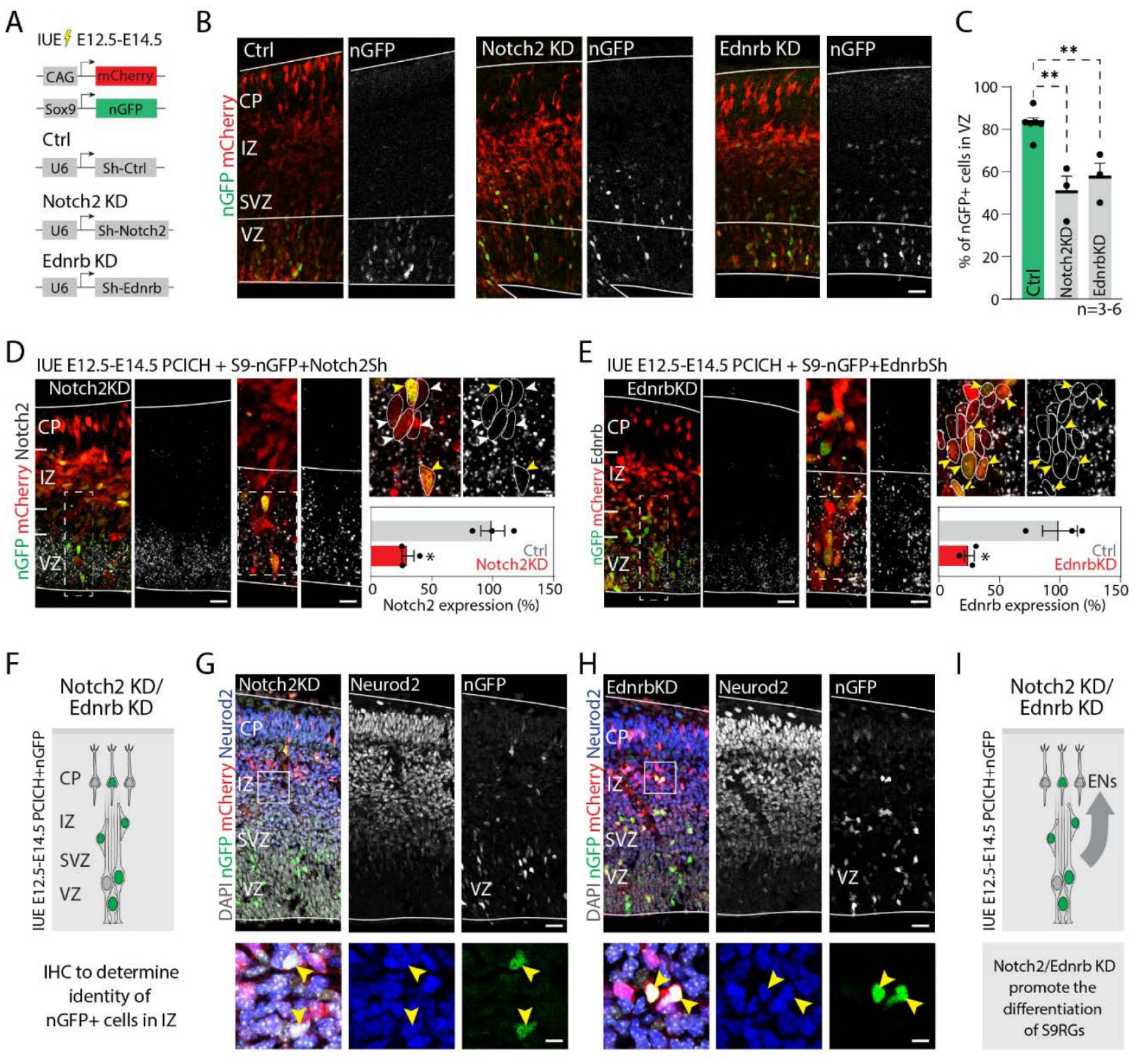
Receptor signaling restrains premature neuronal differentiation. A-B, *Sox9*-nGFP and CAG-mCherry electroporation at E12.5 with scramble, *Notch2* or *Ednrb* shRNA, analyzed 48 h later. C, Fraction of reporter-positive electroporated cells retained in the ventricular zone. D-E, RNAscope validation of *Notch2* and *Ednrb* knockdown. F-H, NEUROD2 staining identifies reporter-positive neurons outside the ventricular zone after knockdown. I, Summary model. Related supplementary figure: S3.

### Selective regulatory accessibility accompanies progenitor maintenance

We examined whether transcriptional differences between S9RGs and comparison RGs were accompanied by changes in chromatin accessibility. For this, we performed ATAC-seq on the sorted populations, using four biological replicates per condition (Fig. 5A). Accessibility at genes used to identify the different cell types was consistent with the enrichment of progenitors in both populations. Moreover, principal component analysis separated S9RGs from comparison RGs (Fig. 5B–C). We identified 3,993 regions with differential accessibility, of which 381 were more accessible and 3,612 were less accessible in S9RGs (FDR < 0.05; Fig. 5D). Genes associated with these regions were enriched in processes related to progenitor proliferation, neurogenesis and neuronal differentiation (Fig. 5E). Interestingly, we found more accessible regions at genes involved in progenitor maintenance, including Notch2 and Ednrb, while regions at neurogenic genes such as Neurod2 and Neurod6 were less accessible (Fig. 5F–G). Thus, despite the overall reduction in accessibility, selected genes involved in progenitor maintenance showed increased accessibility in S9RGs.

**Figure 5.**
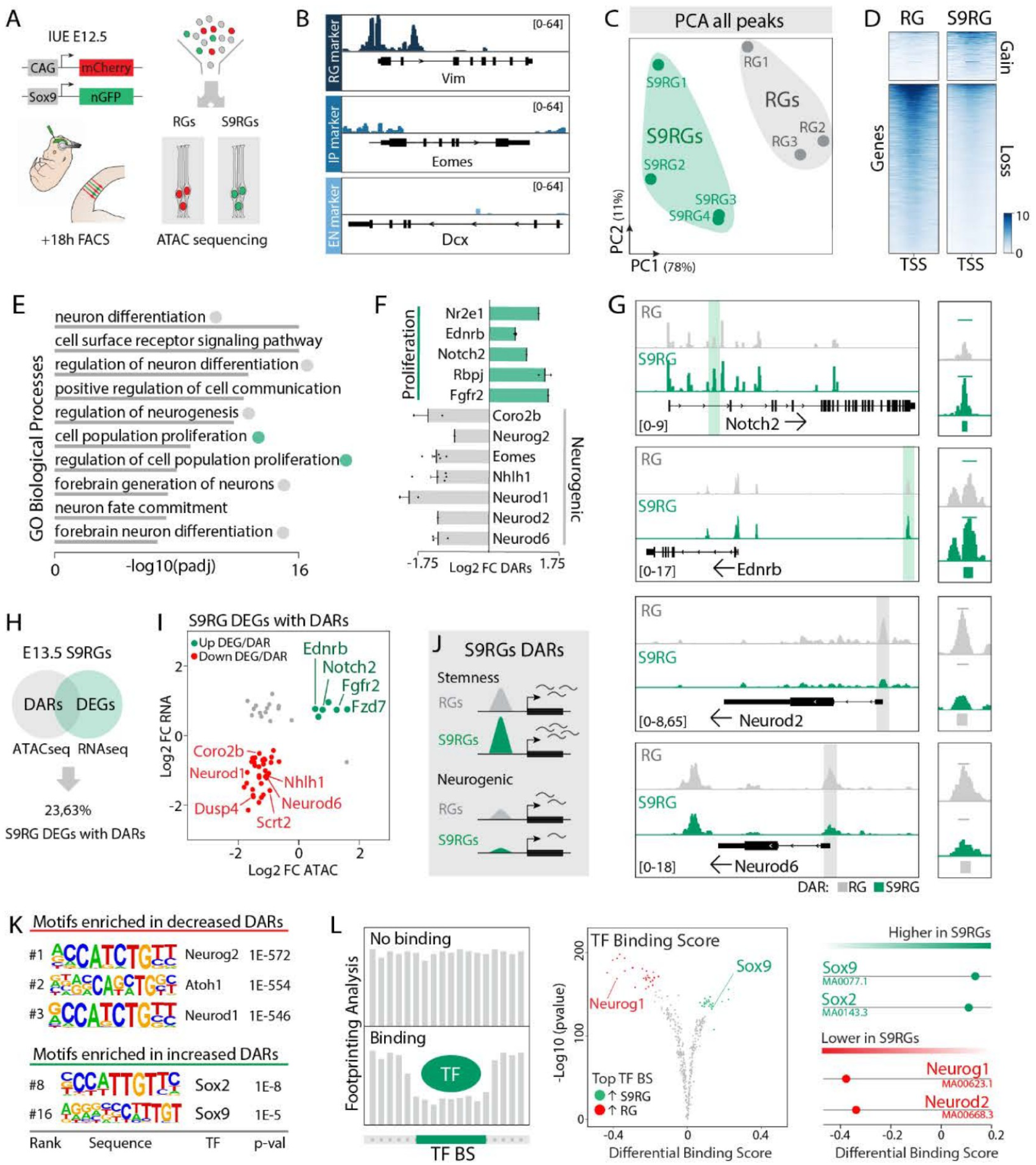
Selective regulatory accessibility accompanies progenitor maintenance. A, ATAC-seq of sorted S9RGs and comparison RGs. B, Accessibility tracks at canonical cell-type markers. C, Principal component analysis; four biological replicates per condition. D, Differential accessibility: 381 regions with increased and 3,612 with reduced accessibility in S9RGs, FDR below 0.05. E, Enrichment of biological processes among associated genes. F-G, Differential regions and tracks at representative maintenance-associated and neurogenic loci. H-I, Integration with differential expression. J, Summary model. K, HOMER motif enrichment in regions gaining or losing accessibility. L, TOBIAS footprint analysis of inferred motif-associated activity. DAR, differentially accessible region. Related supplementary figure: S4.

To relate these chromatin differences to the transcriptional program identified in S9RGs, we compared the chromatin accessibility and gene expression datasets. Some genes showed changes in both expression and chromatin accessibility. For example, Notch2, Ednrb and Fgfr2 were more highly expressed in S9RGs and had regions with increased accessibility, while several neurogenic genes showed the opposite changes (Fig. 5H–I). However, this relationship was not found for all genes. Changes in expression could occur through regions that were already accessible, while some differences in accessibility could appear before changes in gene expression can be detected.

To identify transcription factors potentially associated with these chromatin differences, we analyzed transcription factor binding motifs enriched in differentially accessible regions. Regions that were more accessible in S9RGs were enriched in SOX-family motifs, while less accessible regions were enriched in motifs associated with neurogenic factors, including NEUROG and NEUROD (Fig. 5K). Moreover, ATAC footprint analysis using TOBIAS predicted higher SOX-associated activity and lower activity of proneurogenic factors in S9RGs (Fig. 5L). These results agree with the differences in gene expression and suggest that the chromatin accessibility pattern in S9RGs is consistent with progenitor maintenance and reduced neurogenic differentiation.

### Notch signaling acts upstream of Sox9 in early radial glia

To identify signaling pathways potentially contributing to this program, we analyzed the pathways associated with the top-ranked differentially expressed genes and regions with differential accessibility. Interestingly, Notch was among the main signaling pathways identified in both analyses (Fig. 6A–B). Moreover, S9RGs showed higher overall receptor expression relative to ligand expression, including increased Notch2 and reduced Dll3 and Dll4 expression (Fig. 6C). Together with the premature differentiation observed after Notch2 knockdown, these results suggest that Notch signaling contributes to the molecular properties and maintenance of S9RGs.

**Figure 6.**
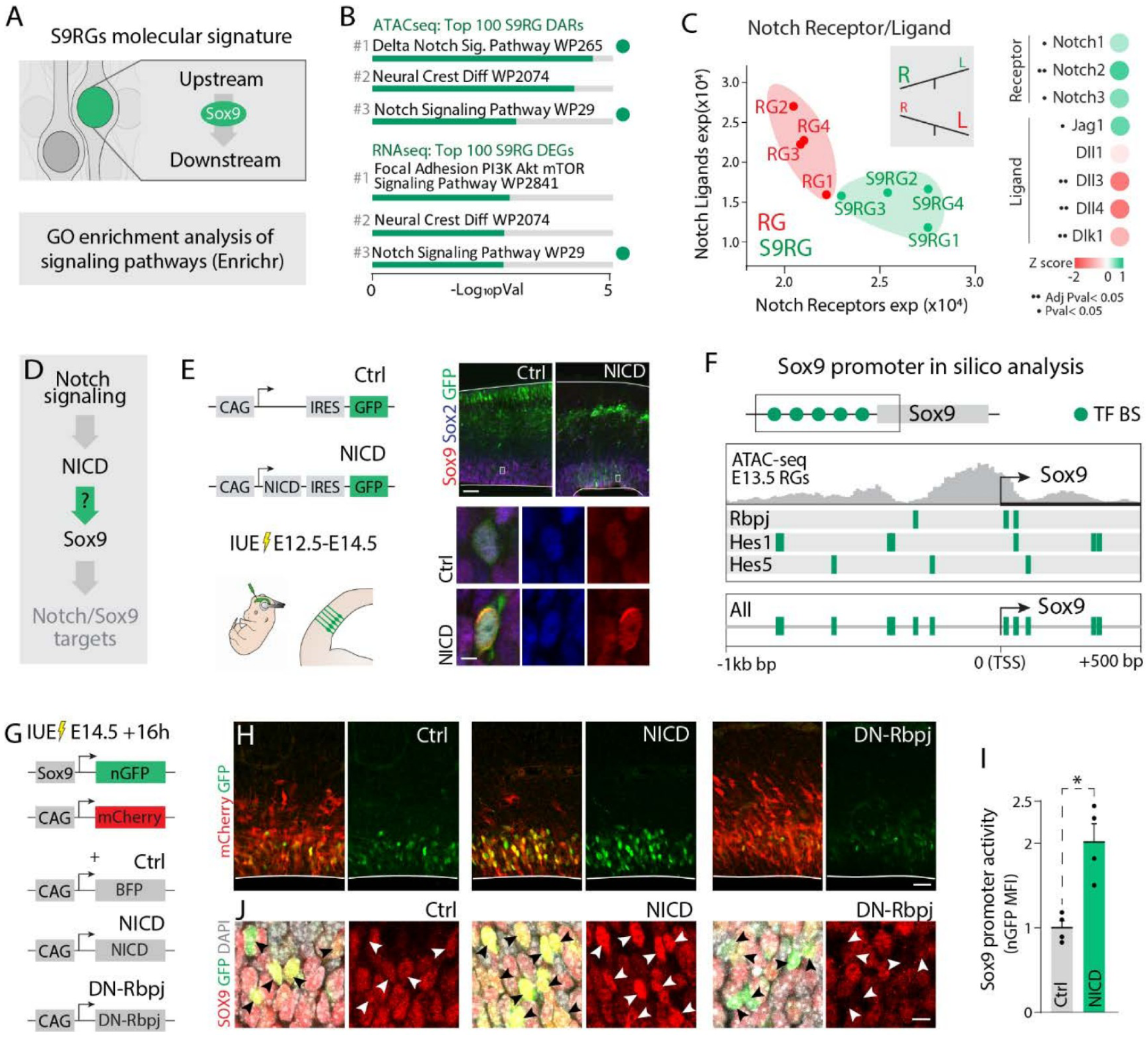
Notch signaling acts upstream of Sox9 in early radial glia. A, Search for upstream signaling pathways using the S9RG molecular signatures. B, Top-ranked pathway terms from the top 100 differential accessibility regions and differential genes; Notch-associated terms are highlighted. C, Combined Notch receptor and ligand expression across RNA-seq samples and individual differential pathway components. D-E, Experimental rationale and NICD expression at E12.5, followed by SOX9 and SOX2 immunostaining at E14.5. F, Predicted Notch-associated transcription factor motifs across the proximal *Sox9* promoter, alongside E13.5 RG ATAC-seq accessibility. G, Reporter perturbation initiated at E14.5 with NICD, dominant-negative RBPJ or control and analyzed 16 h later. H, *Sox9* reporter fluorescence under the three conditions. I, Flow cytometry quantification of reporter intensity after NICD activation. J, Endogenous SOX9 staining under the same reporter perturbation conditions.

Given the regulation of receptor expression by Sox9, we examined whether Notch could also regulate Sox9. For this, we electroporated NICD at E12.5 and analyzed endogenous SOX9 expression at E14.5. NICD increased SOX9 levels in ventricular progenitors (Fig. 6D–E), indicating that activation of Notch signaling can promote Sox9 expression in these cells. To investigate whether this regulation involved Sox9 promoter activity, we examined the proximal Sox9 promoter, which was accessible in E13.5 cortical RGs and contained predicted binding sites for Notch-associated transcriptional regulators, including RBPJ (Fig. 6F). We therefore electroporated the Sox9 promoter reporter together with NICD or dominant-negative RBPJ at E14.5 and analyzed the cells 16 h later. NICD increased reporter fluorescence, while dominant-negative RBPJ reduced it (Fig. 6G–H). We also confirmed the increase in reporter activity after NICD expression by flow cytometry (Fig. 6I). Moreover, endogenous SOX9 levels increased after Notch activation and decreased after its inhibition (Fig. 6J). These results indicate that Notch signaling regulates Sox9 promoter activity and endogenous SOX9 expression in cortical RGs.

### Sox9 connects Notch signaling to progenitor maintenance

To determine whether Notch activation reproduces the expression program identified in S9RGs, we analyzed published RNA-seq data from E13.5 RGs with NICD activation (20). Sox9 was among the genes with increased expression (Fig. 7A–B). Moreover, comparison with our S9RG data showed similar changes among the shared differentially expressed genes. Receptors involved in progenitor maintenance, including Notch2, Ednrb and Ptprz1, were increased, while neurogenic genes such as Neurog1 and Neurod1 were reduced (Fig. 7C–D). These changes were also consistent with those observed after Sox9 overexpression (Fig. 7E–F), suggesting that Notch activation and increased Sox9 expression promote part of the molecular program found in S9RGs. To validate these observations independently, we performed qPCR in CD133-enriched RGs isolated from E13.5 control and Emx1-NICD embryos. These cells showed increased expression of Sox9 and the selected receptors, together with reduced expression of Dll3, Neurog1 and Mfng (Fig. 7G). These results confirmed that Notch activation induces a transcriptional response similar to that found in S9RGs.

**Figure 7.**
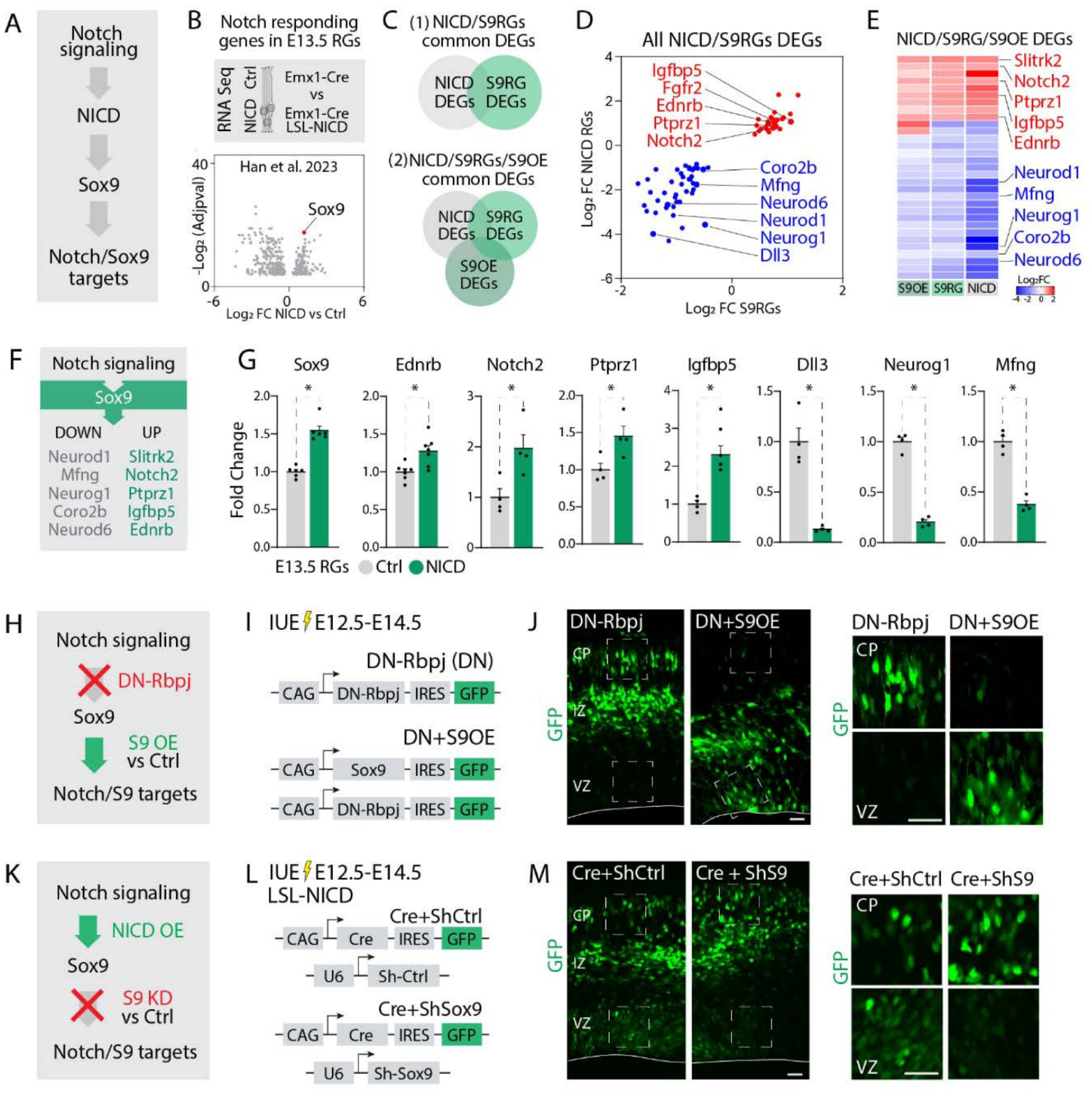
Sox9 connects Notch signaling to progenitor maintenance. A-B, Notch-responsive expression in E13.5 RGs from Han et al. (20), highlighting *Sox9* induction. C-D, Overlap and concordance between NICD-responsive and endogenous S9RG differential genes. E-F, Shared responses across endogenous S9RGs, *Sox9* gain of function and NICD activation, with the inferred regulatory hierarchy. G, Independent qPCR validation in CD133-enriched E13.5 control and Emx1-NICD RGs, four to six biological samples. H-J, *Sox9* overexpression combined with dominant-negative RBPJ after E12.5 electroporation, analyzed at E14.5; representative distributions and ventricular-zone and cortical-plate magnifications. K-M, Mosaic NICD activation in Rosa26-LSL-NICD embryos combined with control or *Sox9* shRNA, analyzed at E14.5. The two perturbation strategies test the functional ordering of Notch and *Sox9* in progenitor retention.

To determine whether Sox9 contributes to Notch-dependent progenitor maintenance, we first electroporated dominant-negative RBPJ alone or together with Sox9 at E12.5 and analyzed the cortex at E14.5. Inhibiting Notch reduced the proportion of labeled cells remaining in the VZ, while Sox9 co-expression counteracted this effect (Fig. 7H–J). Conversely, we examined whether reducing Sox9 expression affected progenitor maintenance following Notch activation. Mosaic NICD activation in Rosa26-LSL-NICD embryos maintained labeled cells in the progenitor compartment. However, simultaneous Sox9 knockdown reduced the proportion of cells remaining in this compartment and increased their contribution to the cortical plate (Fig. 7K–M). These experiments indicate that Sox9 acts downstream of Notch and contributes to its effect on progenitor maintenance. Together with the changes in Sox9 promoter activity, endogenous SOX9 levels and gene expression, these results suggest that Notch maintains early RGs and delays their differentiation in part through Sox9 and the genes it regulates.

### Notch activation produces chromatin accessibility changes similar to those found in S9RGs

Notch activation and Sox9 overexpression produced similar transcriptional changes. We therefore examined whether Notch also modifies chromatin accessibility. For this, we performed ATAC-seq on CD133-enriched RGs isolated from E13.5 control and Emx1-NICD embryos, using three biological replicates per condition (Fig. 8A). Samples separated according to genotype, and we identified 14,438 regions with increased accessibility and 11,239 with reduced accessibility after NICD activation (Fig. 8B–C). Genes associated with more accessible regions were enriched in cell signaling and progenitor proliferation, while those associated with less accessible regions were enriched in neuronal differentiation (Fig. 8D). For example, we found increased accessibility at regions associated with Sox9, Notch1, Notch2 and Hes5. In contrast, selected regions at genes involved in neurogenesis, including Dll1, Eomes, Neurod1 and Neurod6, were less accessible (Fig. 8E–F). These results agree with the increased expression of progenitor-associated genes and reduced expression of neurogenic genes observed after Notch activation.

**Figure 8.**
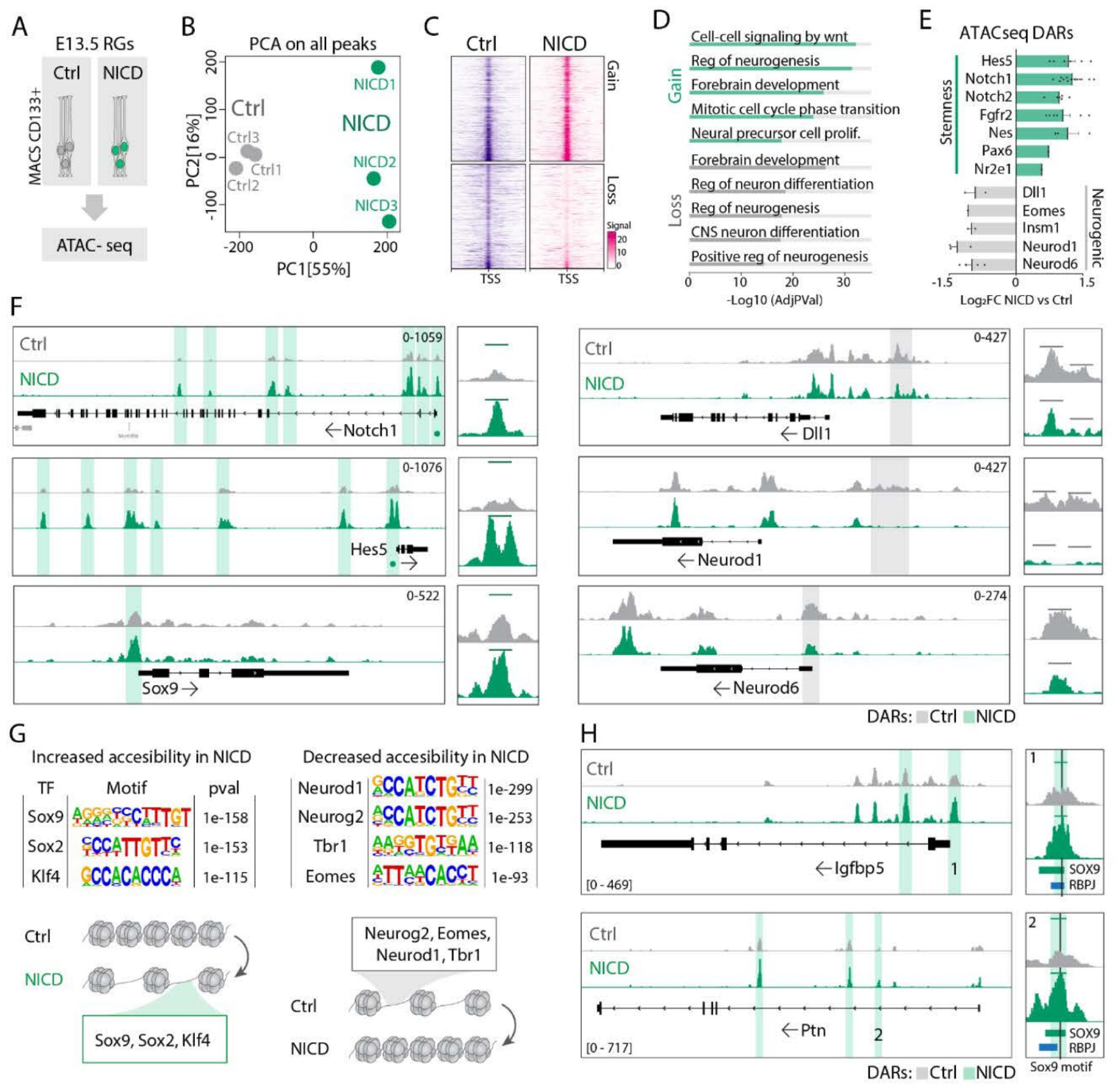
Notch activation converges on the endogenous progenitor regulatory program. A, CD133-based enrichment of E13.5 control and Emx1-NICD RGs for ATAC-seq, three biological replicates per condition. B, Principal component analysis. C, Differential accessibility, including 14,438 increased and 11,239 decreased regions after NICD activation, FDR below 0.05. D, Biological-process enrichment. E-F, Representative maintenance-associated and neurogenic loci and their differential regions. G, Motif enrichment in regions gaining or losing accessibility. H, Selected SOX9 motif-containing regions with public SOX9 and RBPJ ChIP-seq annotations. Related supplementary figure: S4.

We also examined the transcription factor binding motifs present in these regions. Regions with increased accessibility after NICD activation were enriched in progenitor-associated motifs, including those of the SOX family, while less accessible regions were enriched in proneurogenic motifs (Fig. 8G). Moreover, some regions containing SOX9 motifs overlapped SOX9 or RBPJ binding sites reported in public datasets (Fig. 8H). To determine whether similar chromatin changes were present in S9RGs, we compared the genes associated with differential accessibility in both datasets. Interestingly, 89.2% of the genes associated with differential regions in S9RGs were also associated with differential accessibility after NICD activation (fig. S4A). Among these shared genes, changes in expression generally followed the same direction in S9RGs and NICD-activated RGs (fig. S4B). Moreover, selected genes involved in progenitor maintenance and neurogenesis showed similar changes in both expression and accessibility (fig. S4C). Together, these results suggest that Notch activation reproduces part of the transcriptional and chromatin accessibility differences found in S9RGs, particularly at genes involved in progenitor maintenance and neurogenesis.

### Differences in neuronal identity genes are already present before RGs differentiate

We examined whether the molecular differences in S9RGs extend to genes involved in neuronal identity, beyond those regulating progenitor maintenance and neurogenesis. For this, we analyzed chromatin accessibility at selected genes involved in lower- and upper-layer neuronal specification (Fig. 9A). We found that several lower-layer-associated genes, including Tbr1, Bcl11b and Sox5, had regions with reduced accessibility in S9RGs (Fig. 9B–C). In contrast, the upper-layer-associated genes examined maintained their accessibility. Comparison with our gene expression data showed that some of the lower-layer-associated genes also showed reduced expression in S9RGs. For example, Tbr1 had both reduced accessibility and lower expression, which we confirmed by qPCR (Fig. 9D–E). These results indicate that the molecular differences between the two progenitor populations are not limited to genes regulating proliferation and differentiation, but also include genes involved in the specification of the neurons they later generate.

**Figure 9.**
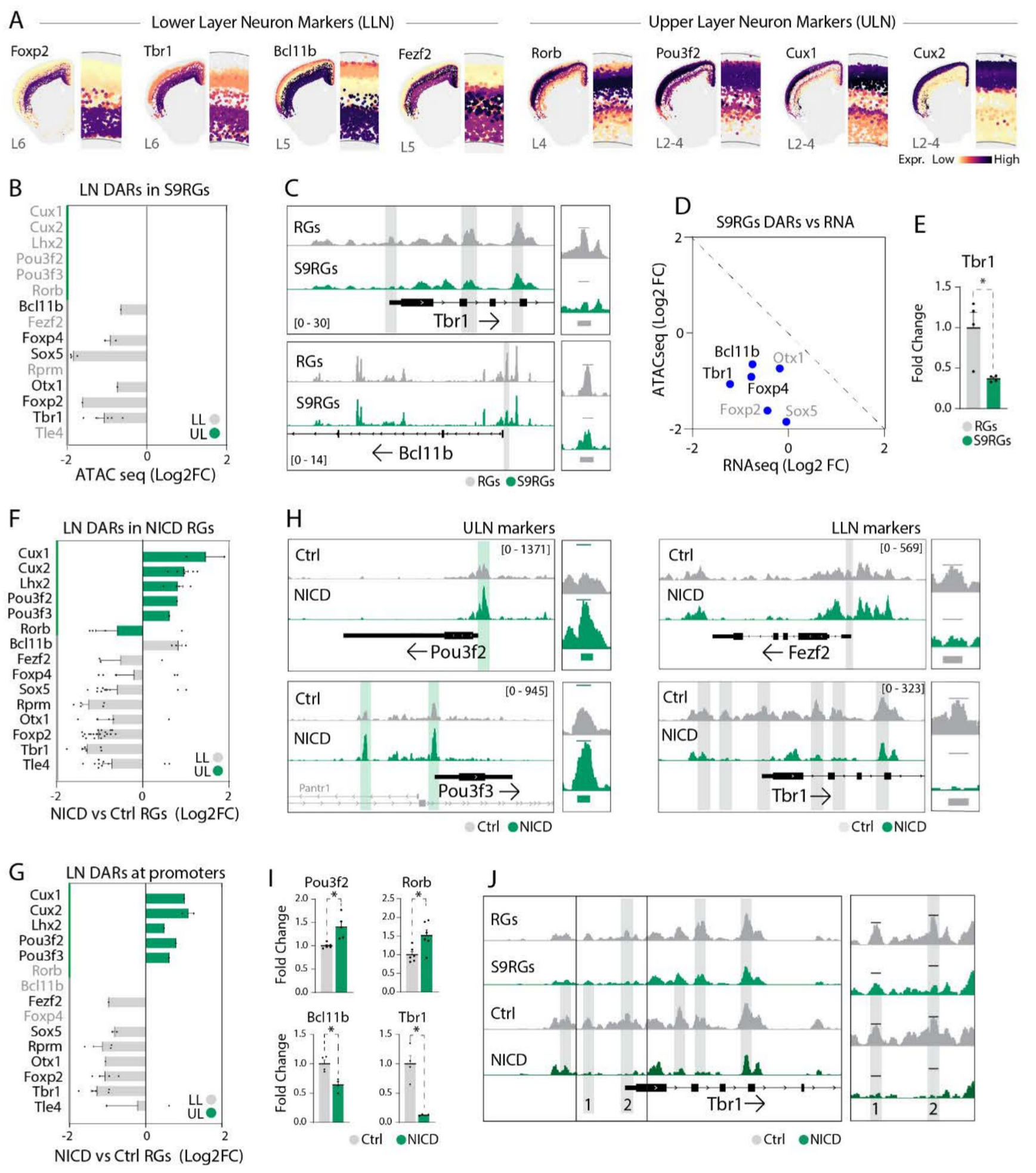
Neuronal identity programs diverge before radial glia differentiate. A, Adult reference expression maps of layer-associated genes from the Allen Brain Cell Atlas. B-E, S9RG accessibility and expression at selected deep-layer-associated loci. F-I, Accessibility and representative qPCR responses after NICD activation. J, Representative *Tbr1* accessibility comparison. Accessibility and expression profiles relate early progenitor regulation to later neuronal identity programs. Related supplementary figure: S5.

We also examined whether Notch activation produced similar changes. NICD overexpression reduced accessibility at selected lower-layer-associated genes. Moreover, it increased accessibility at several upper-layer-associated genes, including Pou3f2, Pou3f3, Lhx2 and Cux2 (Fig. 9F–G). We examined these changes at individual loci and confirmed selected expression differences by qPCR, including reduced Tbr1 and Bcl11b and increased Pou3f2 (Fig. 9H–I). Interestingly, comparison of the Tbr1 locus in S9RGs and NICD-activated RGs showed regions with accessibility changes in the same direction (Fig. 9J). We further examined Cux2, which showed increased accessibility at several regions together with higher expression in CD133-enriched RGs after Notch activation. Using RNAscope combined with PAX6 and TBR2 staining, we detected the increased expression in VZ progenitors (fig. S5). These results indicate that the change in Cux2 expression was already present before neuronal differentiation. Together, these results show that S9RGs have reduced accessibility at selected lower-layer-associated genes while maintaining accessibility at the upper-layer-associated genes examined. Notch activation produces similar changes and also increases accessibility at several upper-layer-associated genes. These early molecular differences could contribute to the preferential generation of upper-layer neurons by SOX9-high progenitors, together with their delayed neurogenesis.

### Components of the S9RG molecular program are present in other cortical progenitor populations

To examine the S9RG program in independently profiled mouse progenitors, we used the developing cortical atlas from Di Bella et al. (2), selected the cells annotated as RGs and scored their expression of the 103 genes enriched in S9RGs. Cells with high scores were already present at early neurogenic stages, and the program was more represented at later ages in this analysis (fig. S6A–F). Moreover, electroporation of the Sox9 reporter at successive embryonic ages showed an increasing proportion of reporter-positive cells in the VZ (fig. S6G). To examine whether these differences were also associated with endogenous Sox9 expression, we compared early RGs with different Sox9 levels, independently of the signature score. Interestingly, Sox9-high cells showed gene expression changes similar to those found in our bulk comparison of S9RGs and comparison RGs (fig. S6H–I). These results indicate that the molecular differences identified using the reporter are also present among cortical progenitors in an independent dataset.

We also examined the expression of this program in progenitors from the developing ferret cortex. In the dataset from Del-Valle-Anton et al. (25), alpha and beta RGs both include ventricular and outer populations. We found that 36.9% of the S9RG-enriched genes examined were differentially expressed between alpha and beta RGs, with most expressed at higher levels in alpha RGs (Fig. 10A). These included NOTCH2, EDNRB, PTPRZ1, VCAM1 and MLC1. Their preferential expression in alpha RGs was also observed in expression maps and violin plots (Fig. 10B). Thus, several receptors and cell-surface components enriched in mouse S9RGs are also preferentially expressed in a progenitor population in the ferret cortex.

**Figure 10.**
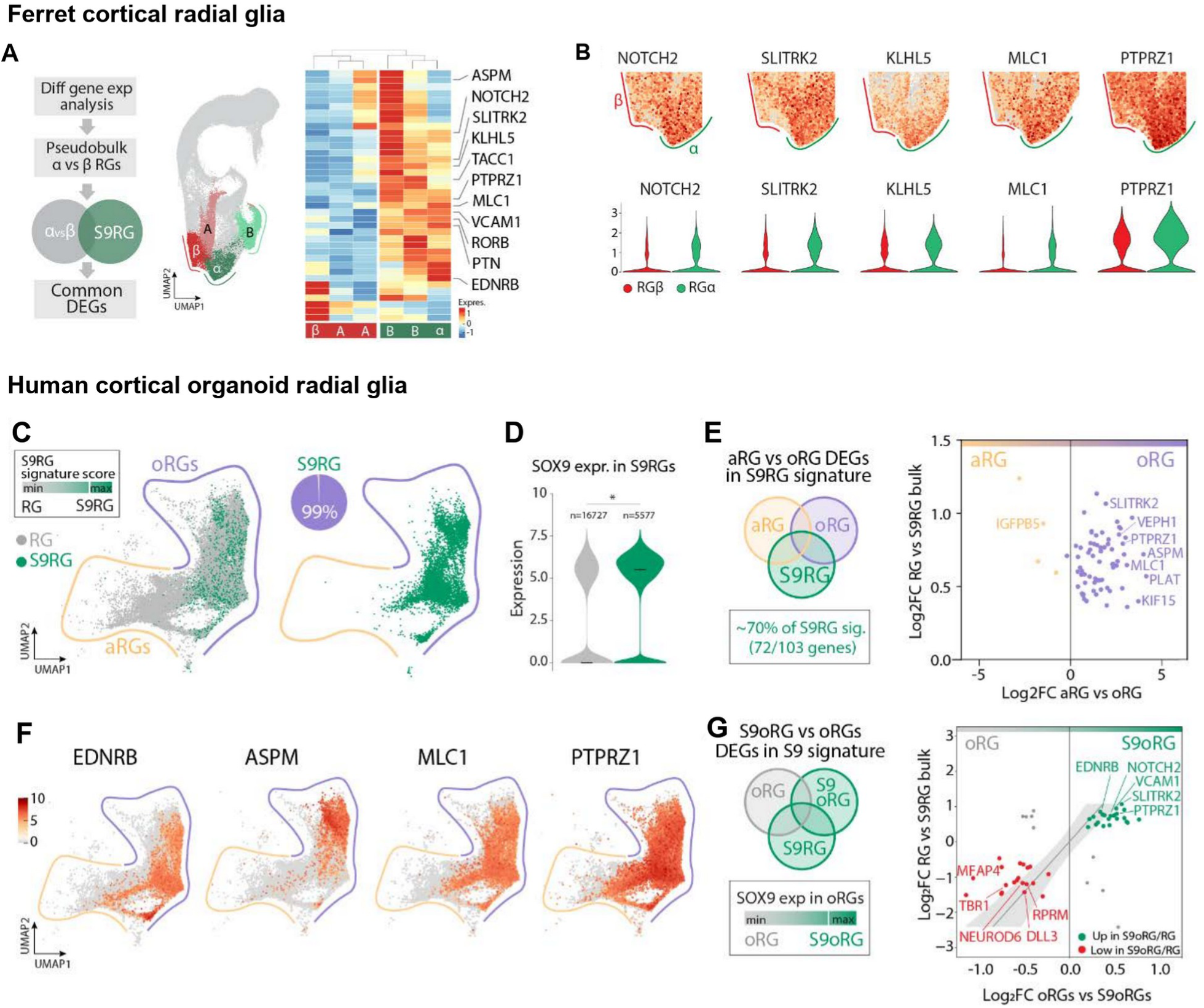
The SOX9-associated program extends across cortical progenitor architectures. A-B, Ferret cortical single-cell analysis from Del-Valle-Anton et al. (25). A, Differential expression between alpha and beta RGs and representation of mouse S9RG-enriched genes. B, Selected gene expression across alpha and beta RGs. C-G, Three-month human cortical organoid analysis from Uzquiano et al. (26). C, S9RG signature scoring; approximately 99% of high-scoring cells are annotated as outer RGs, rather than 99% of all outer RGs expressing the signature. D, SOX9 expression in high-scoring and remaining RGs. E, Differential signature genes between apical and outer RGs. F, Representative enriched genes. G, Concordance between SOX9-high versus remaining outer RGs and the mouse signature. Related supplementary figures: S6, S7.

We extended this comparison to RGs from three-month human cortical organoids using the dataset from Uzquiano et al. (26). Approximately 99% of the cells classified as S9RG-like according to the signature score belonged to the outer RG population (Fig. 10C). Moreover, these cells expressed higher levels of SOX9 (Fig. 10D), indicating that the relationship between SOX9 expression and the S9RG molecular program was also present in human organoid progenitors. Comparison of apical and outer RGs showed that approximately 70% of the signature genes were differentially expressed, and 96% of these were higher in outer RGs (Fig. 10E–F). To examine whether these differences were also present within the outer RG population, we compared outer RGs according to their SOX9 expression levels. SOX9-high outer RGs also showed expression differences similar to those found in mouse S9RGs (Fig. 10G). We also compared one-to-one orthologs across the three datasets and identified signaling and cell-surface components shared by mouse S9RGs, ferret alpha RGs and human organoid outer RGs (fig. S7). Together, these analyses show that components of the S9RG molecular program are present in progenitor populations from different species, despite differences in their position and organization within the developing cortex.

## Discussion

Our previous work showed that early RGs with higher SOX9 expression remain as progenitors for longer and preferentially generate upper-layer neurons (15). Here, we have identified a broader molecular program involving extracellular signaling, progenitor maintenance and neurogenesis that could contribute to these behaviors. Among the genes enriched in S9RGs, we identified several signaling receptors, and reducing Notch2 or Ednrb expression promotes their differentiation. Moreover, we show that Notch regulates Sox9 expression and that Sox9 contributes to progenitor maintenance downstream of this pathway. Interestingly, S9RGs also show differences in chromatin accessibility at genes involved in neuronal identity. These findings suggest that the mechanisms maintaining early progenitors could also influence the neurons they later generate.

Interestingly, S9RGs delay neurogenesis but already express some genes characteristic of later RGs. This suggests that their temporal molecular properties can change while they remain as progenitors. Previous studies showed that temporal gene expression programs are transmitted from RGs to their neuronal descendants and can progress independently of cell division (1, 11). Our results suggest that these temporal properties also differ among progenitors present at the same embryonic age. Moreover, increasing Sox9 expression promotes some of these later molecular features without significantly increasing the differentiation score. Our lineage-tracing experiments show that S9RGs preferentially contribute to ULNs. In our previous work, the few neurons found in the lower layers of the Hes5 lineage expressed SATB2 but not CTIP2, suggesting that the contribution of these progenitors also differs in projection neuron identity (15). The molecular differences identified here could therefore contribute to their neuronal preference, together with their delayed neurogenesis.

The increased expression of receptors in S9RGs could help explain how these cells remain as progenitors for longer. We show that Sox9 regulates the expression of several of these receptors and that reducing Notch2 or Ednrb promotes neuronal differentiation. These results indicate that at least part of the molecular program identified in S9RGs contributes to their maintenance. Interestingly, the expression of genes involved in extracellular signaling also increases in later RGs (1). Differences in receptor expression could therefore change how progenitors respond to their environment as development progresses. For example, NTF3 released by differentiating cortical neurons can regulate progenitor behavior and the transition from lower- to upper-layer neuronal production (28). Although we have not identified the signals acting through the receptors enriched in S9RGs, their increased expression could allow these cells to respond differently from neighboring progenitors to the signals present in the developing cortex.

Our experiments suggest that Notch maintains early cortical progenitors in part through Sox9. We found that Sox9 expression increases after Notch activation. Moreover, Sox9 overexpression allows more cells to remain in the VZ when Notch is inhibited, while Sox9 knockdown reduces the proportion of cells remaining as progenitors after Notch activation. A role for Sox9 downstream of Notch was previously described in embryonic stem cell-derived cultures, where it contributes to stem cell maintenance and astroglial differentiation (55). Here, we show that Sox9 also contributes to the response to Notch in early cortical RGs and regulates genes enriched in S9RGs.

Among them, Notch2 is particularly interesting because its expression increases after Sox9 overexpression, and its reduction promotes neuronal differentiation. In agreement with these results, overexpression of activated NOTCH2 was previously shown to maintain proliferation and inhibit differentiation of cerebellar granule neuron progenitors (56). Thus, by increasing Notch2 expression, Sox9 could help progenitors maintain their response to Notch signals. This raises the possibility that Notch and Sox9 reinforce each other, although this feedback remains to be tested. It would also be interesting to study how this regulation depends on the duration and dynamics of Notch activity. Oscillatory expression of Notch effectors contributes to neural progenitor maintenance (17), and these dynamics are not reproduced by our NICD experiments.

We also found differences in chromatin accessibility that agreed with the changes in gene expression in S9RGs. For example, Notch2 and Ednrb had regions with increased accessibility, while several neurogenic genes showed reduced expression and accessibility. Notch activation produced similar changes in the expression and accessibility of selected genes involved in progenitor maintenance and neurogenesis. These results suggest that Notch could contribute to the molecular differences found in S9RGs. Previous work on SOX2 also showed a relationship between chromatin regulation and NSC maintenance. Sox2 deletion disrupted enhancer–promoter interactions and reduced gene expression, and restoring Socs3 expression rescued the self-renewal defect (57).

Interestingly, in S9RGs we also found differences at genes involved in neuronal identity. Several lower-layer-associated regions were less accessible, while the upper-layer-associated regions examined remained accessible. Notch activation produced similar changes and also increased accessibility at some upper-layer-associated genes. These differences could influence the neuronal programs expressed when progenitors later differentiate. In agreement with this possibility, cortical progenitors already express transcripts associated with neuronal identity before the corresponding proteins are detected (22). Moreover, Ring1B regulates the termination of subcerebral projection neuron production, while DOT1L contributes to the early regulation of upper-layer-associated genes in progenitors (23, 24). Our results suggest that the signals maintaining early RGs could also participate in the regulation of neuronal identity genes. At later stages, Notch also promotes chromatin changes that allow progenitors to acquire gliogenic competence (21), indicating that this pathway can regulate both progenitor maintenance and their ability to generate different cell types.

These findings could help explain why early cortical progenitors make different contributions to neuronal populations. Fate-mapping and clonal studies have identified progenitors with preferential contributions to particular neuronal subtypes, although individual clones also show considerable variation (3, 6–10). Our results suggest that delayed neurogenesis and early molecular differences could contribute together to this behavior. Their preferential contribution to ULNs could therefore depend both on when they generate neurons and on the molecular properties of the progenitors from which these neurons arise. These properties may also influence how they respond to the changing cortical environment. This possibility is consistent with early differences among progenitors, while allowing their neuronal contribution to be modified as development progresses.

Interestingly, components of the S9RG molecular program were also found in ferret alpha RGs and human cortical organoid outer RGs (25, 26). In ferret, alpha RGs include both ventricular and outer progenitors and are enriched in proliferation-associated genes (25). Thus, the similarities with S9RGs could reflect shared mechanisms of progenitor maintenance rather than their position in the cortex. In human organoids, we also found SOX9-associated expression differences within the outer RG population, indicating that the relationship was not simply due to comparing apical and outer progenitors. Previous studies showed that human oRGs express genes involved in extracellular interactions and self-renewal, including PTPRZ1, which is also enriched in S9RGs (29, 30). Moreover, live imaging of human bRGs showed Notch activation in the daughter cell that remained as a progenitor (58). Together, these observations support the importance of signaling pathways for progenitor maintenance in different cortical populations. Although our comparisons do not establish that the shared genes have the same function in each population, they suggest that part of this program is already present in early mouse ventricular RGs. Changes in the regulation of these signaling components could also be relevant to cortical evolution. For example, human-specific NOTCH2NL genes promote progenitor maintenance through Notch regulation, while a human-gained enhancer regulates FGFR2 expression and progenitor proliferation (31–33). Our findings raise the possibility that differences in when and where these shared genes are expressed contribute to the diversity of cortical progenitor behavior.

Our findings show that the differences among early cortical RGs extend beyond SOX9 expression. S9RGs have a molecular program involving receptors, regulators of progenitor maintenance and differences in chromatin accessibility at genes associated with neuronal identity. We show that Notch and Sox9 regulate components of this program and that selected receptors contribute to maintaining these cells as progenitors. These molecular properties could help explain their preferential contribution to ULNs, together with their delayed neurogenesis. Thus, maintaining progenitors for later development could influence both the number and the types of neurons they generate. The presence of components of this program in ferret and human progenitors further suggests that changes in its regulation could have contributed to the diversification of progenitor behavior during cortical evolution.

## Materials and methods

### Animals and in vivo manipulations

We used C57BL/6J, Ai9 (Jackson Laboratory, 007909; 62), Emx1-Cre (005628; 63) and Rosa26-LSL-NICD (008159; 64) mice. All lines were maintained on a C57BL/6 background. Animals were housed under a 12 h light–dark cycle with food and water available ad libitum. The day of vaginal plug detection was considered E0.5, and embryos of both sexes were used. Emx1-NICD embryos were obtained by crossing Emx1-Cre with Rosa26-LSL-NICD mice, as previously described (20). All animal experiments performed at the University of Valencia were approved by its Ethics Committee and the corresponding regional authority.

The constructs used in this work, their origin, references and shRNA target sequences are listed in Supplementary Table S1. We generated Sox9-nGFP by cloning a proximal fragment of the mouse Sox9 promoter, approximately −2 kb to +100 bp relative to the transcription start site, into a vector containing a minimal promoter and nuclear GFP. We also used this fragment to drive Cre expression and permanently label the progeny of Sox9-reporter-positive cells in Ai9 embryos. Constructs generated in our laboratory were verified by whole-plasmid sequencing. To activate NICD in a mosaic manner, we electroporated Cre into Rosa26-LSL-NICD embryos.

In utero electroporation was performed as described in our previous work (15, 59). Pregnant females were anesthetized with isoflurane (2.5% induction, 1–1.5% maintenance), and buprenorphine was administered at 0.1 mg/kg. A plasmid solution containing 0.75–1.5 µg/µl of each construct was injected into one embryonic lateral ventricle. Electroporation was performed using a BTX830 electroporator, with the electrodes oriented towards the lateral cortex. Five pulses were applied at 30 V for 50 ms in E12.5 embryos or at 35 V for 70 ms in E13.5 embryos, with a 950 ms interval between pulses. Embryos were collected at the ages indicated in each experiment.

### Histology and image analysis

Brain tissue was processed for immunohistochemistry as previously described (15). Embryonic brains were fixed in 4% paraformaldehyde for 4–8 h at 4 °C. For vibratome sectioning, brains were embedded in 4% low-melting-point agarose and cut coronally at 100 µm. For cryostat sectioning, brains were cryoprotected in 30% sucrose, embedded in OCT and cut at 12–14 µm. Sections were incubated in blocking buffer containing 10% horse serum and 0.2% Triton X-100 in PBS for 1 h. Primary antibodies were diluted in blocking buffer and incubated overnight at 4 °C. The following day, sections were washed and incubated with fluorophore-conjugated secondary antibodies for 1 h at room temperature. DAPI was used to label nuclei. Antibodies are listed in Supplementary Table S2.

RNAscope was performed using the Multiplex Fluorescent Detection Kit v2 (Advanced Cell Diagnostics, 323110), following the manufacturer’s instructions with modifications to allow subsequent immunostaining. Cryostat sections were post-fixed, dehydrated and treated with hydrogen peroxide. We omitted antigen retrieval and reduced the protease treatment to 5 min to improve antibody staining. Sections were then incubated with the probes for 2 h at 40 °C, followed by signal amplification and tyramide detection. We used probes for Ednrb (473801), Notch2 (425161), Ptprz1 (460991) and Vcam1 (438641). After RNAscope, sections were processed for immunohistochemistry as described above.

Images were acquired using Olympus FV10i and Zeiss LSM 980 confocal microscopes. We quantified the distribution of labeled cells and fluorescence intensity in the cortical compartments indicated in each experiment. For RNAscope, individual electroporated cell bodies in the VZ were outlined in Fiji and saved in the ROI Manager. Puncta within each cell were counted using Find Maxima, with prominence values between 6 and 12 depending on the probe. The same settings were used for all samples within each comparison.

### Radial glial isolation and RNA analysis

Electroporated cortical regions were dissected and the meninges removed. Cortices were incubated with trypsin at 37 °C and mechanically dissociated in buffer containing DNase. To select RGs, we used the surface markers CD133 and CD24, as previously described (**60**). Cells were incubated for 30 min on ice with CD133-BV421 (BioLegend, 141213; 1:400) and CD24-PerCP-Cy5.5 (BD, 562360; 1:400). DAPI (0.1 µg/ml) was used to identify dead cells. We first excluded debris, doublets and dead cells, and then selected electroporated CD133-positive cells with low or no CD24 expression. The fluorescent reporters allowed us to separate the populations of interest using a FACSAria Fusion, keeping samples at 4 °C. For RNA analysis, we collected at least 20,000–30,000 cells per sample, pooling two or three embryos when needed. For ATAC-seq of S9RGs and comparison RGs, we collected approximately 30,000–50,000 cells per sample. To isolate RGs from control and Emx1-NICD embryos, cortices were dissociated with papain and CD133-positive cells were selected using magnetic microbeads (Miltenyi, 130-092-564), MS columns and an OctoMACS separator.

RNA was purified using the RNeasy Plus Micro Kit according to the manufacturer’s instructions. We used the SuperScript IV Single Cell/Low Input cDNA PreAmp Kit to reverse-transcribe the RNA and preamplify the cDNA for 16 cycles. To measure gene expression, we performed qPCR in a StepOnePlus Real-Time PCR System using TB Green Premix Ex Taq. We first checked the efficiency of each primer pair using standard curves and its specificity by melting curve analysis. Gene expression was normalized to Gapdh and calculated using the 2^−ΔΔCt method (61).

To compare the transcriptomes of S9RGs and comparison RGs, we sequenced four independently sorted samples per population. For Sox9 overexpression, we used six biological replicates per condition obtained from at least three litters. Libraries were prepared using the Novogene NGS RNA Library Prep Set and sequenced on an Illumina NovaSeq X Plus, using paired-end 150 bp reads and targeting approximately 15 Gb of raw data per sample. We filtered the reads with fastp, aligned them using HISAT2 v2.0.5 and obtained gene-level counts with featureCounts v1.5.0-p3 (34–36). Differential expression was analyzed using DESeq2 with Benjamini–Hochberg correction. Genes with an adjusted P ≤ 0.05 were considered differentially expressed (37). To study the effects of Notch activation, we also analyzed the published RNA-seq data from E13.5 NICD-activated RGs obtained by Han et al. (20).

### Chromatin accessibility analysis

To study chromatin accessibility in these progenitor populations, we performed ATAC-seq (38) using the Zymo-Seq ATAC Library Kit (D5458), following the manufacturer’s instructions. We used four biological replicates per population for S9RGs and comparison RGs, and three per genotype for control and Emx1-NICD RGs. Libraries were size-selected between approximately 150 and 500 bp and sequenced using paired-end 150 bp reads. For the S9RG comparison, we targeted 120 Gb of raw data per sample. For the NICD experiment, libraries were prepared from 50,000 CD133-enriched cells per sample and sequenced to approximately 30 million reads per sample.

Bioinformatic analysis was performed using nf-core/atacseq v2.1.2 (39). Reads were aligned to the mouse reference genome GRCm38 with BWA, and mitochondrial reads were excluded (40). We used sambamba v0.8.2 to retain fragments shorter than 100 bp for the analysis of nucleosome-free regions (41). Peaks were called using MACS2 v2.1.1 in paired-end mode, keeping duplicate reads (42). To compare accessibility between conditions, we used DiffBind with DESeq2. Regions with FDR < 0.05 were considered significantly different and annotated with ChIPseeker v1.38.0, using a 3 kb window around the transcription start site for promoter-proximal assignment (43, 44). Coverage tracks were generated with deepTools and visualized in IGV (45, 46). For comparison between datasets, we used the genes associated with differential regions in each experiment and identified those common to both analyses.

To identify transcription factor binding motifs enriched in regions with differential accessibility, we used HOMER (47). We also performed footprint analysis with TOBIAS, using the HOCOMOCO v14 motif collection (48, 49). Functional enrichment was analyzed with Enrichr (50). To compare accessible regions with previously reported SOX9 and RBPJ binding sites, we obtained public binding annotations from ChIP-Atlas (51). In situ hybridization images used to examine gene expression in the developing cortex were obtained from GenePaint (52).

To look for binding sites for Notch-associated transcriptional regulators in the Sox9 promoter, we analyzed the region from −1 kb to +500 bp relative to the transcription start site. We used the Search Motif Tool from the Eukaryotic Promoter Database (https://epd.expasy.org), selecting predicted sites with P < 0.001. These sites were plotted together with the ATAC-seq signal from E13.5 cortical RGs.

### Single-cell dataset analysis

We analyzed published scRNA-seq data from mouse cortex (2) and human cortical organoids (26), obtained from the Single Cell Portal under accession numbers SCP1290 and SCP1756, respectively. Ferret data were provided by the laboratory of Víctor Borrell (25). We used the original cell annotations to select early and mid-neurogenic mouse RGs, ferret alpha and beta RGs at E34 and P1, and RGs from three-month human organoids. To examine the expression of the S9RG program in these populations, we used AUCell (53) with the 103 genes upregulated in our S9RG samples (adjusted P < 0.05; log2 fold change > 0.35). Cells with scores in the upper quartile were considered S9RG-like. We also selected SOX9-high cells according to their normalized SOX9 expression, using the upper quartile independently of the signature score.

To compare gene expression between species, we identified high-confidence one-to-one orthologs using Ensembl through biomaRt (54). We added the counts from cells belonging to the same RG population within each biological sample and used DESeq2 for differential expression analysis. In ferret, we compared alpha with beta RGs. In human organoids, we compared apical with outer RGs, and SOX9-high outer RGs with the rest of the outer RG population. We then compared the changes in expression of the corresponding orthologs with those found in our mouse S9RG analysis. To study temporal properties and differentiation, we used the gene sets and scores described by Telley et al. (1). We first checked these scores in independently isolated CD133-enriched RG and neuron-enriched populations from E13 and E16 cortices, with four samples per group.

### Statistical analysis

Statistical analyses were performed using GraphPad Prism 9. We first tested data normality using the Shapiro–Wilk test. For comparisons between two groups, we used Student’s t test or the Mann–Whitney test. For three or more groups, we used ANOVA followed by Tukey’s multiple comparisons or the Kruskal–Wallis test. Multiple-testing corrections for sequencing data are described in the corresponding sections above. Data are shown as mean ± SEM unless otherwise indicated. Rescue panels and selected morphological observations were assessed qualitatively.

## Data and code availability

We will deposit the RNA-seq and ATAC-seq data generated in this work in GEO and make them publicly available upon publication. The published mouse cortical and human organoid datasets used in our analyses are available in the Single Cell Portal under accession numbers SCP1290 and SCP1756, respectively. Ferret data were provided by the laboratory of Víctor Borrell (25).

## Acknowledgments

We thank Dr. Isabel Fariñas and Dr. Isabel Martínez-Garay for access to reagents and equipment, and Dr. Víctor Borrell for providing the ferret single-cell dataset. We also thank the staff of the University of Valencia Research Support Services (SCSIE), particularly the animal facility, microscopy and flow cytometry services, for their assistance.

## Supplementary materials

**Supplementary Figure S1.**
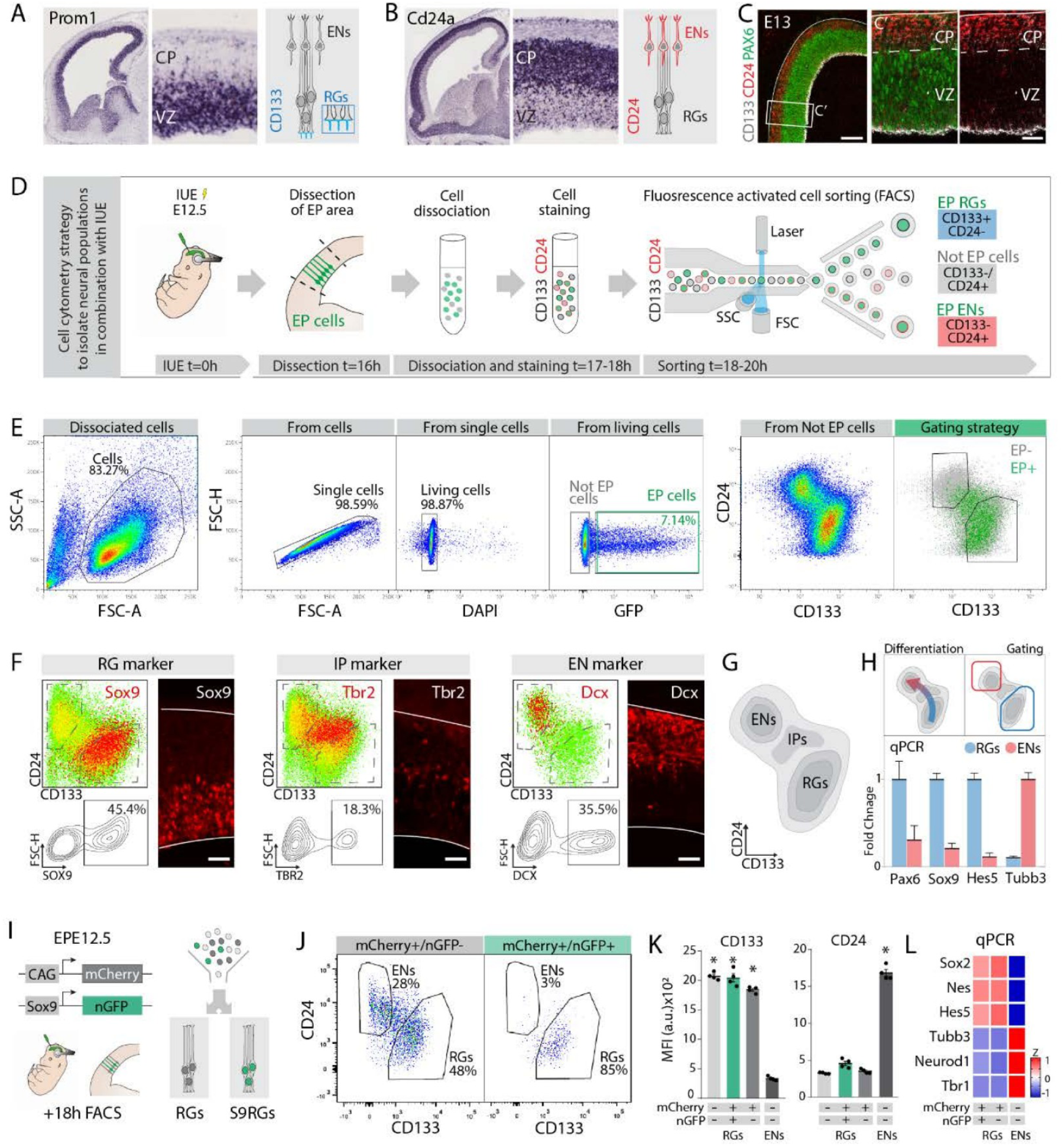
Reporter-defined populations retain radial glial identity during isolation. Related to Figures 1 and 2. Validates population identity and purification. A-C, Prom1/CD133 and Cd24a/CD24 reference expression and tissue staining. D-E, Dissociation and FACS selection of viable singlets and electroporated cells. F-G, Intracellular SOX9, TBR2 and DCX validation of CD133/CD24 compartments. H, Differentiation axis and qPCR marker validation. I-J, Reporter-based separation of S9RGs and comparison RGs within the enriched progenitor compartment. K-L, Surface marker levels and qPCR validation of cell identity.

**Supplementary Figure S2.**
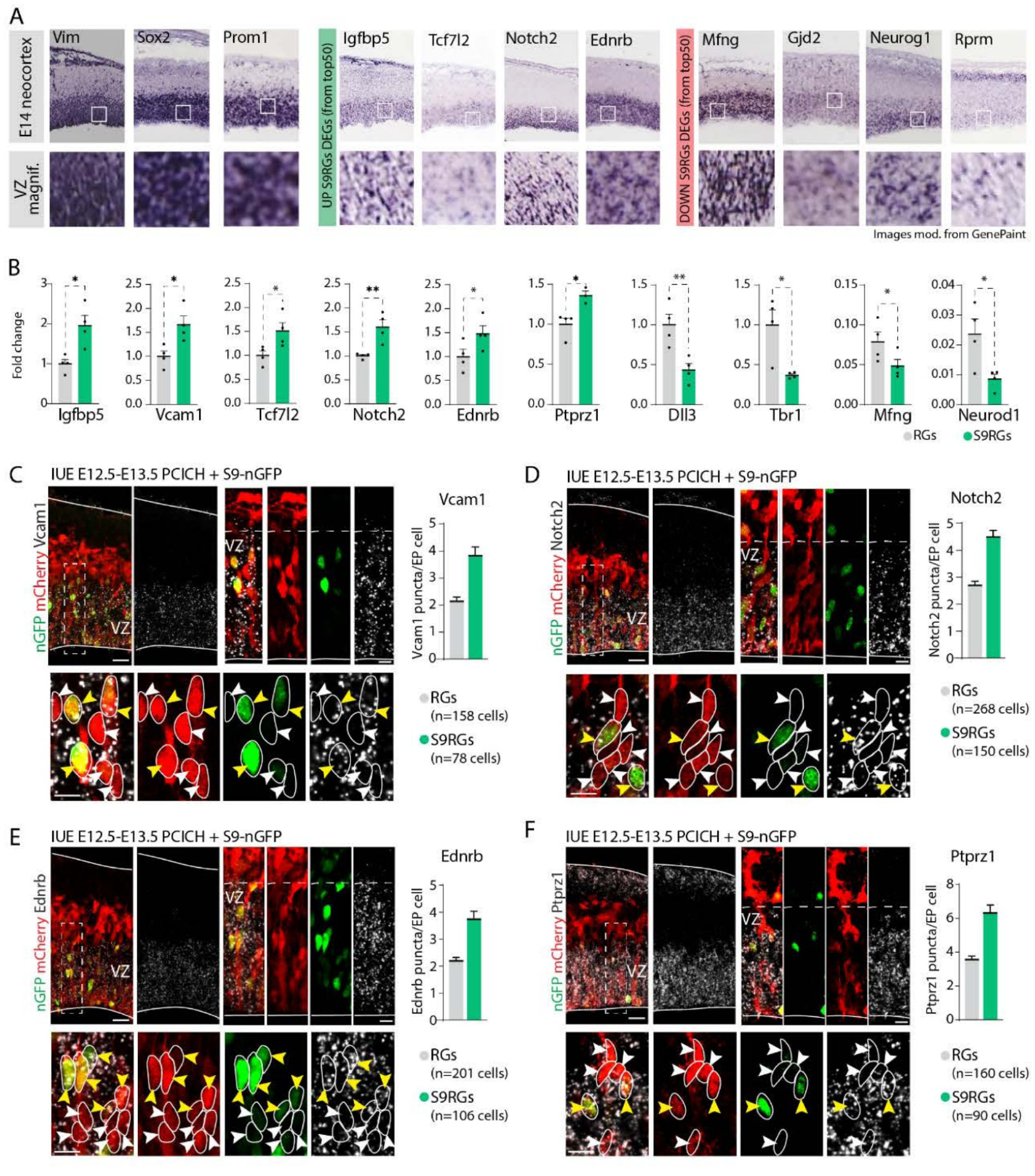
The expression program is detected within ventricular-zone progenitors. Related to Figure 2. Validates differential gene expression in tissue. A, GenePaint in situ hybridization examples illustrating cortical and ventricular-zone expression patterns. B, qPCR validation of representative S9RG differential genes, with four biological samples. C-F, RNAscope for *Vcam1*, *Notch2*, *Ednrb* and *Ptprz1* after electroporation of *Sox9*-nGFP and CAG-mCherry at E12.5, analyzed at E13.5. Yellow arrowheads identify examples of double-positive S9RGs and white arrowheads identify comparison cells. Puncta per electroporated cell are shown alongside the corresponding image series; numbers of analyzed cells are shown in the plots.

**Supplementary Figure S3.**
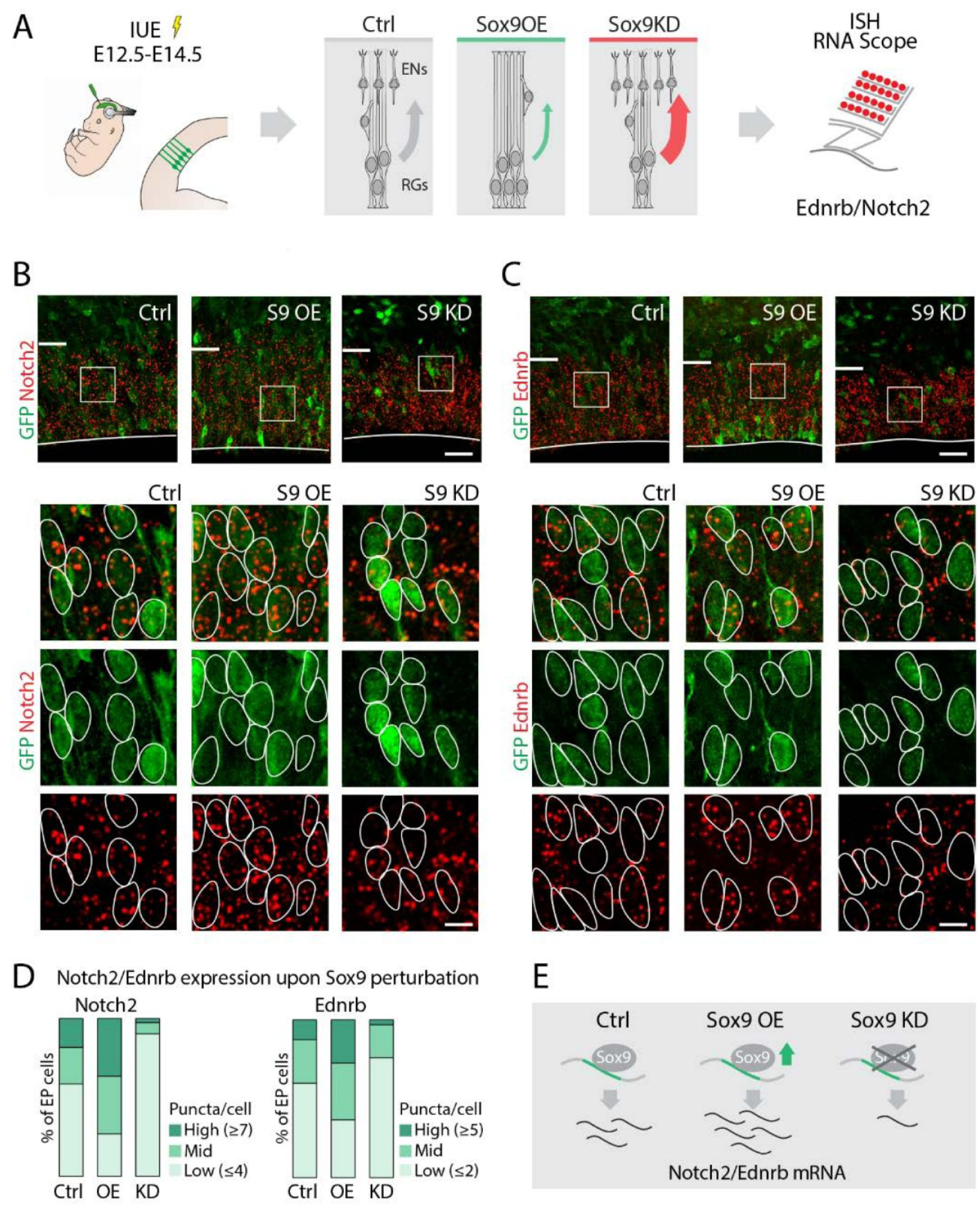
Endogenous Sox9 levels regulate the receptor program. Related to Figures 3 and 4. Tests reciprocal transcript responses to *Sox9* gain and loss of function. A, Electroporation followed by RNAscope. B-C, *Notch2* and *Ednrb* expression at E14.5 after E12.5 control, *Sox9* overexpression or *Sox9* knockdown electroporation. D, Distribution of cells among high, intermediate and low puncta categories. E, Summary of the inferred regulatory relationship. Bars show the fraction of electroporated cells in each transcript-abundance category.

**Supplementary Figure S4.**
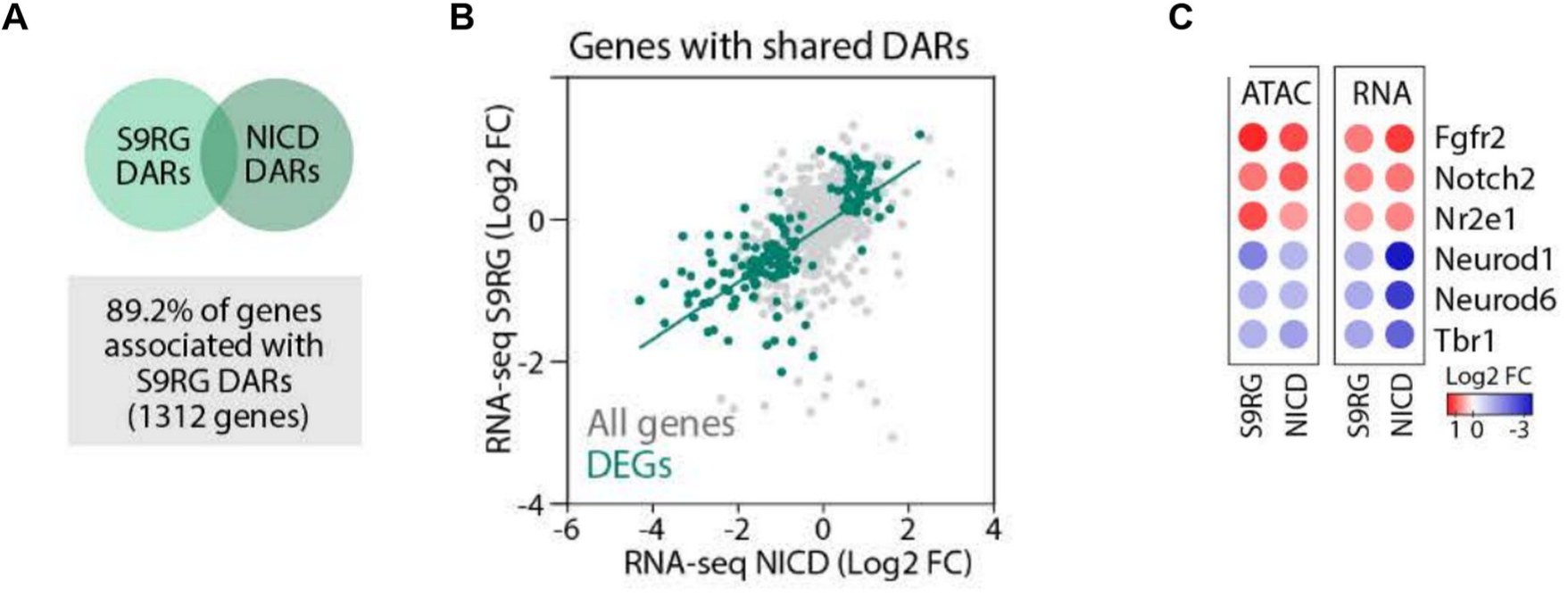
Notch activation converges with the S9RG chromatin program. Related to Figures 5 and 8. A, Overlap among genes associated with differential accessibility in endogenous S9RGs and NICD-activated RGs; 89.2% refers to genes associated with S9RG differential accessibility regions. B, Concordance of transcriptional changes among genes associated with differential accessibility in both comparisons. C, Representative expression and accessibility responses at maintenance- and neuronal-associated genes.

**Supplementary Figure S5.**
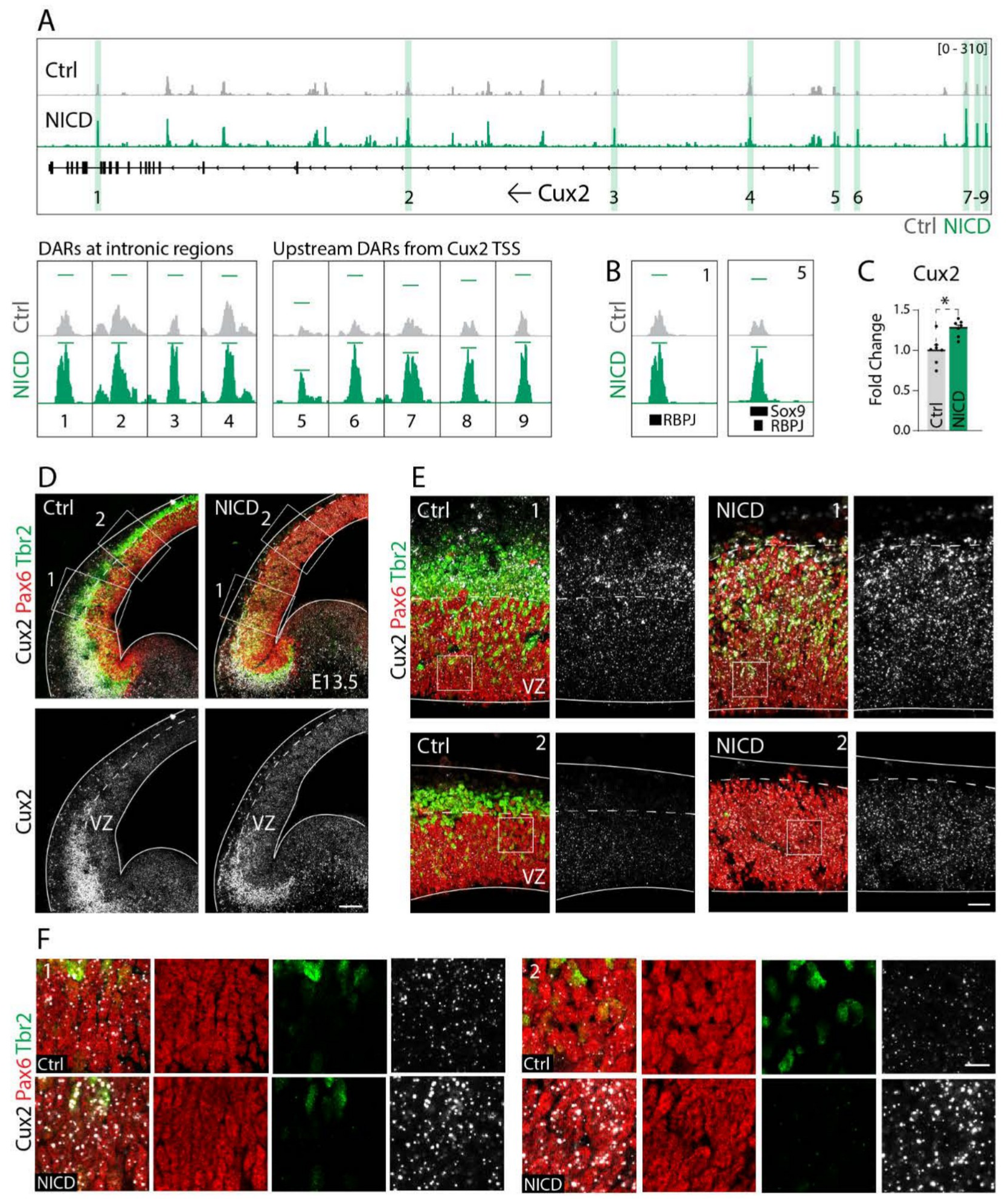
Notch induces Cux2 expression before neuronal differentiation. Related to Figure 9. Validates the NICD-associated *Cux2* response. A-B, NICD-responsive accessibility at the *Cux2* locus and public SOX9/RBPJ ChIP-seq annotations. C, qPCR in CD133-enriched E13.5 control and Emx1-NICD RGs. D-F, RNAscope for *Cux2* combined with PAX6 and TBR2 staining in control and NICD cortex. NICD activation induces *Cux2* expression in cortical progenitors.

**Supplementary Figure S6.**
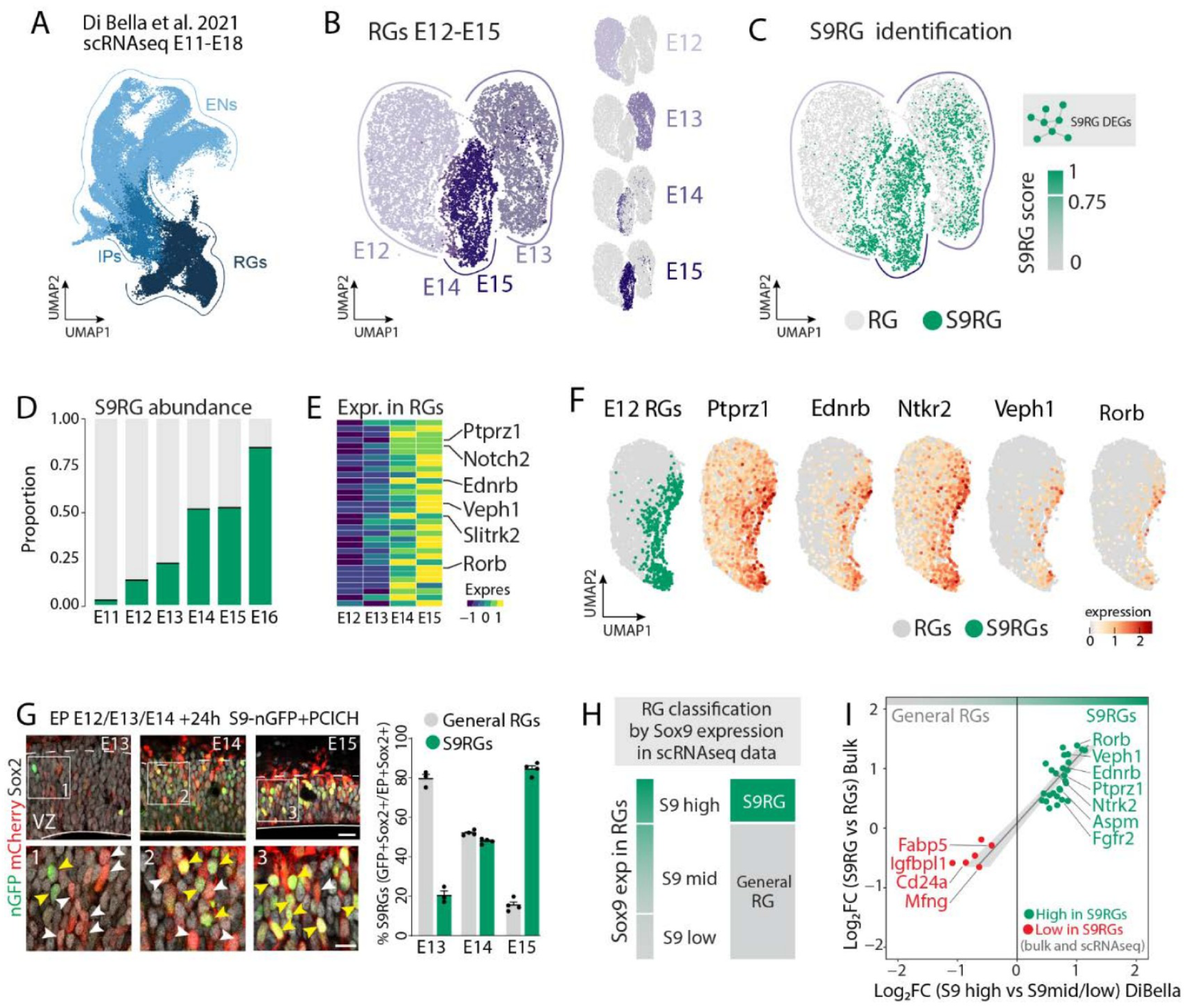
Independent mouse data recover SOX9-associated progenitor heterogeneity. Related to Figure 10. Tests the signature independently in mouse before comparative analysis. A-F, Analysis of the mouse cortical atlas of Di Bella et al. (2), retaining excitatory-lineage and RG annotations, with AUCell signature scores and selected marker expression. G, Reporter-positive fractions in the ventricular zone after electroporation at successive embryonic stages and collection 24 h later. H-I, Comparison of early *Sox9*-high RGs with remaining RGs and correspondence with the bulk S9RG signature. Signature scores and SOX9 expression classes were defined using upper-quartile thresholds.

**Supplementary Figure S7.**
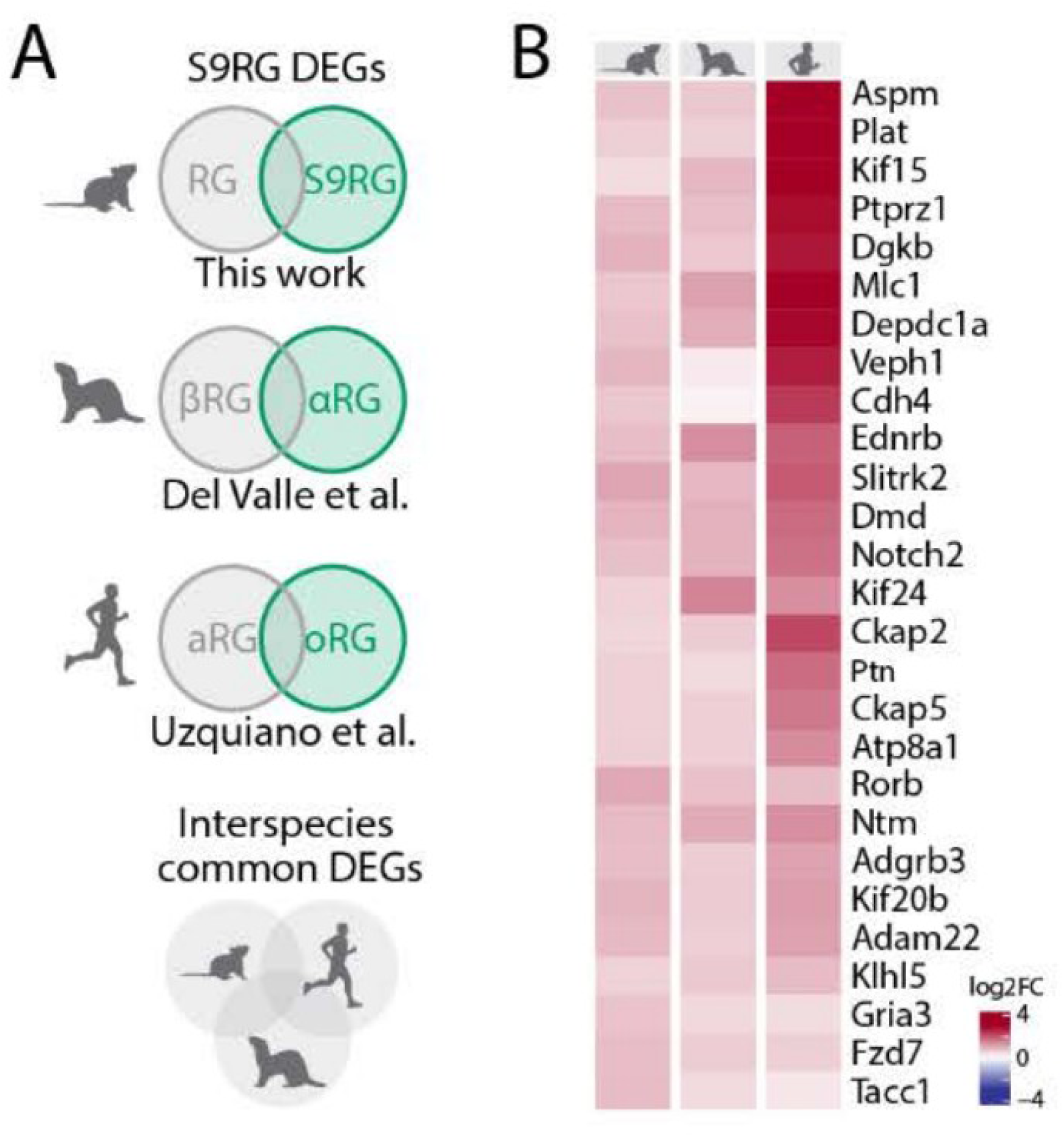
Orthologous components connect progenitor programs across species. Related to Figure 10. Summarizes shared orthologous components. A-B, Cross-species comparison and heatmap of S9RG-enriched genes shared among mouse S9RGs, ferret alpha RGs and human organoid outer RGs. Shared orthologues identify recurrent components of the progenitor program across species.

**Supplementary Table S1.**
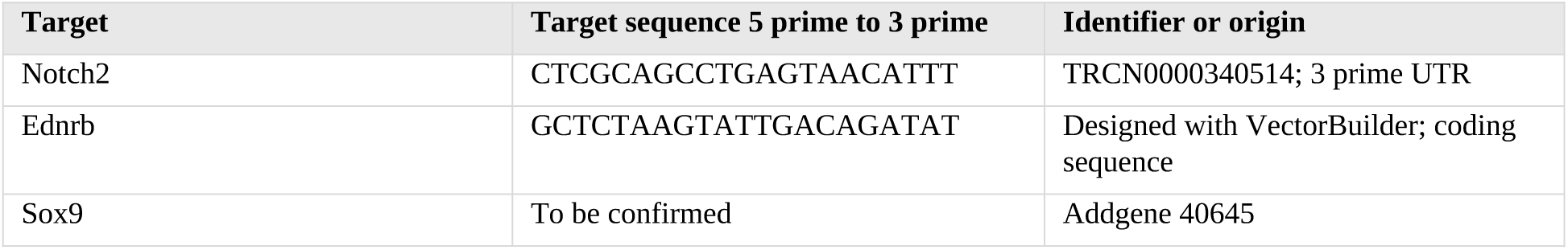
shRNA target sequences.

**Supplementary Table S2.**
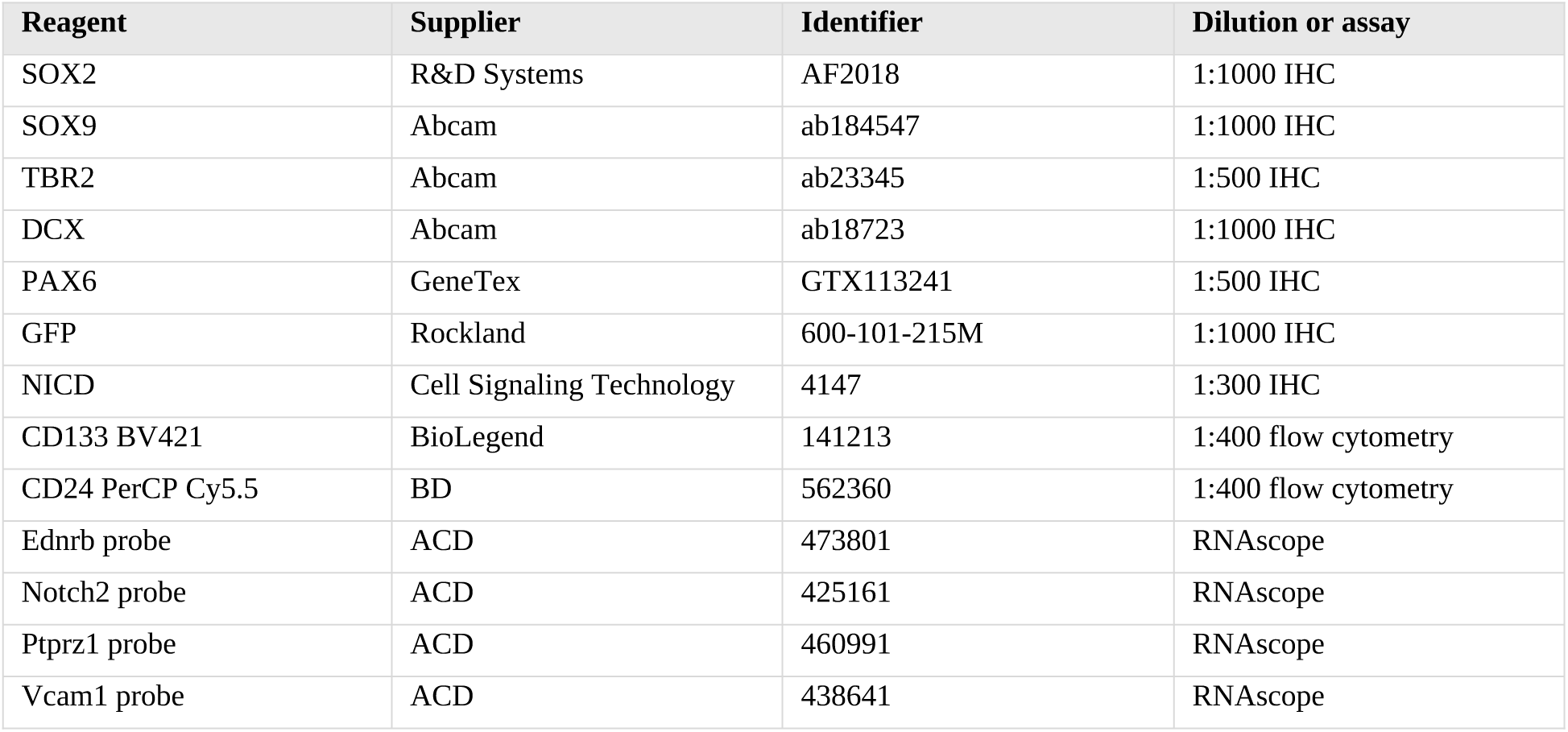
Antibodies and probes.

| **Reagent** | **Supplier** | **Identifier** | **Dilution or assay** |
|---|---|---|---|
| SOX2 | R&D Systems | AF2018 | 1:1000 IHC |
| SOX9 | Abcam | ab184547 | 1:1000 IHC |
| TBR2 | Abcam | ab23345 | 1:500 IHC |
| DCX | Abcam | ab18723 | 1:1000 IHC |
| PAX6 | GeneTex | GTX113241 | 1:500 IHC |
| GFP | Rockland | 600-101-215M | 1:1000 IHC |
| NICD | Cell Signaling Technology | 4147 | 1:300 IHC |
| CD133 BV421 | BioLegend | 141213 | 1:400 flow cytometry |
| CD24 PerCP Cy5.5 | BD | 562360 | 1:400 flow cytometry |
| Ednrb probe | ACD | 473801 | RNAscope |
| Notch2 probe | ACD | 425161 | RNAscope |
| Ptprz1 probe | ACD | 460991 | RNAscope |
| Vcam1 probe | ACD | 438641 | RNAscope |

